# Reprogramming Factors Activate a Non-Canonical Oxidative Resilience Pathway That Can Rejuvenate RPEs and Restore Vision

**DOI:** 10.1101/2025.08.30.673239

**Authors:** Yuancheng Ryan Lu, James C. Cameron, Yan Hu, Han Shen, Shintaro Shirahama, Alexander Tyshkovskiy, Zhaoyi Chen, Jiahe Ai, Daniel Y. Zhu, Margarete M. Karg, Lindsey A. Chew, George W. Bell, Siddhartha G. Jena, Yue He, Philip Seifert, Daisy Y. Shu, Mohamed A. EI-Brolosy, Qiannuo Lou, Bohan Zhang, Anna M. Puszynska, Xiaojie Qiu, Xiao Tian, Meredith Gregory-Ksander, Vadim N. Gladyshev, David A. Sinclair, Magali Saint-Geniez, Jason D. Buenrostro, Catherine Bowes Rickman, Bruce R. Ksander, Jonathan S. Weissman

## Abstract

Oct4, Sox2, and Klf4 (OSK) Yamanaka factors induce pluripotency and reverse age-related epigenetic changes, yet the mechanisms by which they promote rejuvenation remain poorly explored. Oxidative stress contributes to CNS aging and retinal pigmented epithelium (RPE) degeneration in age-related macular degeneration. We find that OSK expression in RPE restores retinal structure and visual function in aged mice and promotes oxidative resilience through a non-canonical, Tet2-independent pathway. Integrative functional genomics identifies GSTA4, a detoxifying enzyme that clears the lipid peroxidation byproduct 4-HNE, as a necessary and sufficient OSK effector. Dynamic GSTA4 regulation by OSK recapitulates a stem cell derived stress resilience program. GSTA4 overexpression alone enhances mitochondrial resilience, rejuvenates the aged RPE transcriptome, and reverses visual decline. GSTA4 is consistently upregulated across diverse lifespan-extending interventions suggesting a broader pro-longevity role. These findings uncover a previously unrecognized protective axis driven by Yamanaka factors that circumvents reprogramming, providing therapeutic insights for age-related diseases.

**HIGHLIGHTS:**

- OSK–GSTA4 provides a dynamic, Tet2-independent stress-resilience axis.
- Functional genomics pinpoints GSTA4 as a direct downstream effector activated by OSK.
- RPE aging involves progressive accumulation of 4-HNE that can be detoxified by GSTA4.
- Enhancing GSTA4 rejuvenates RPE cells, restores vision and is associated with lifespan-extending interventions.

## INTRODUCTION

Aging is a complex and multifaceted degenerative process involving a variety of molecular mechanisms such as epigenetic alterations, telomere attrition, dysregulated nutrient sensing and oxidative stress. Among them, oxidative stress can impact nearly all major hallmarks of aging, most notably genomic instability, mitochondrial dysfunction, cellular senescence, and stem cell exhaustion ^1^, by both impairing protein function ^2^ and generating toxic byproducts. Among all tissues, the role of oxidative stress in driving degeneration is perhaps most clearly established in the retina, an extension of the central nervous system. Oxidative stress emerges early in both the brain and retina and progressively worsens with age ^3^, fueled by environmental exposures, metabolic byproducts, and declining mitochondrial function. This cumulative stress ultimately disrupts cellular integrity and undermines tissue homeostasis.

Age-related macular degeneration (AMD), the leading cause of irreversible vision loss affecting over 200 million people worldwide ^4^, is a prime example of oxidative stress-driven pathology. Dry form of AMD, which accounts for 90% of cases, is driven by degeneration of the retinal pigment epithelium (RPE), a layer highly vulnerable to oxidative damage from chronic light exposure and bisretinoid lipofuscin buildup which elevates reactive oxygen species (ROS) over time^5^. The causal role of ROS in AMD is supported by the AREDS2 study, where antioxidant supplementation slowed disease progression ^6^. The two recently approved therapies for dry AMD treatment provide only modest benefit, likely reflecting the fact that they target components of the alternative complement pathway, a cascade that is activated after oxidative stress has already injured the retina, rather than addressing that upstream damage directly ^7^. This highlights an unmet need to identify novel pathways that enhance oxidative resilience and counteract ROS-induced damage.

Over the past decade, partial epigenetic reprogramming through transient expression of all or subsets of the Yamanaka factors (Oct4, Sox2, Klf4, and c-Myc, aka OSKM) has emerged as a promising strategy to restore youthful tissue function *in vivo* ^8–13^. Dual AAV-mediated delivery of OSK without the c-Myc oncogene has been shown to rejuvenate post-mitotic retinal ganglion cells (RGCs), promoting axon regeneration and restoring vision in either glaucomatous or aged mice, with long-term expression via AAV2 proving safe for up to 18 months ^9,14^. This system also has demonstrated efficacy in multiple sclerosis mouse models ^15^. Beyond the eye, AAV-OSK induces epigenetic rejuvenation in kidney and muscle ^16^, and reproducibly contributes to lifespan extension in aged wild-type mice ^17,18^. Notably, it has also shown promise in a non-human primate model of non-arteritic anterior ischemic optic neuropathy, a common optic neuropathy ^19^.

Despite its therapeutic potential, the mechanisms through which OSK(M) exerts functional rejuvenation remain poorly defined. A few mediators of partial epigenetic reprogramming, including Tet1/2^9^ and Top2a^20^, have been identified, that facilitate chromatin and DNA modifications in cooperation with OSK. However, the broader network of OSK downstream effectors, those functional units that directly carry out the biological effects, remain less well explored. Additionally, toxicity of prolonged OSKM expression has been reported in liver, intestine^21^ and a few brain cell types such as astrocyte and microglia ^12^, which could limit broad application of partial epigenetic reprogramming. A better understanding of how OSK(M) acts, and what its downstream effectors are, could enable more specific and safer approaches of rejuvenation, bypassing the associated safety concerns of prolonged expression ^21,22^.

Here we explore the effect of OSK partial reprogramming in RPE cells, which operate under high oxidative load offering a robust model for probing how rejuvenation programs confer resistance to oxidative challenges. Enabled by a functional genomics approach, our study uncovers a rejuvenation axis involving GSTA4 activation, that bypasses reprogramming-induced dedifferentiation. This pathway acts in a dynamic regulation manner instead of through Tet2 to protect against oxidative damage, potentially opening new therapeutic avenues for aging and age-related diseases.

## RESULTS

### OSK Protects the RPE from Oxidative Stress and Age-related Degeneration

Although ROS buildup has been documented in aging human RPE ^23^, it has not been directly measured in mouse RPE. To address this gap, we performed ROS staining and observed a significant, age-related increase in ROS in murine RPE (**Figure 1A**-**1B**). These findings validate aged mice as a model that recapitulates human RPE oxidative damage and support their use for testing novel therapeutic interventions.

**Figure 1.**
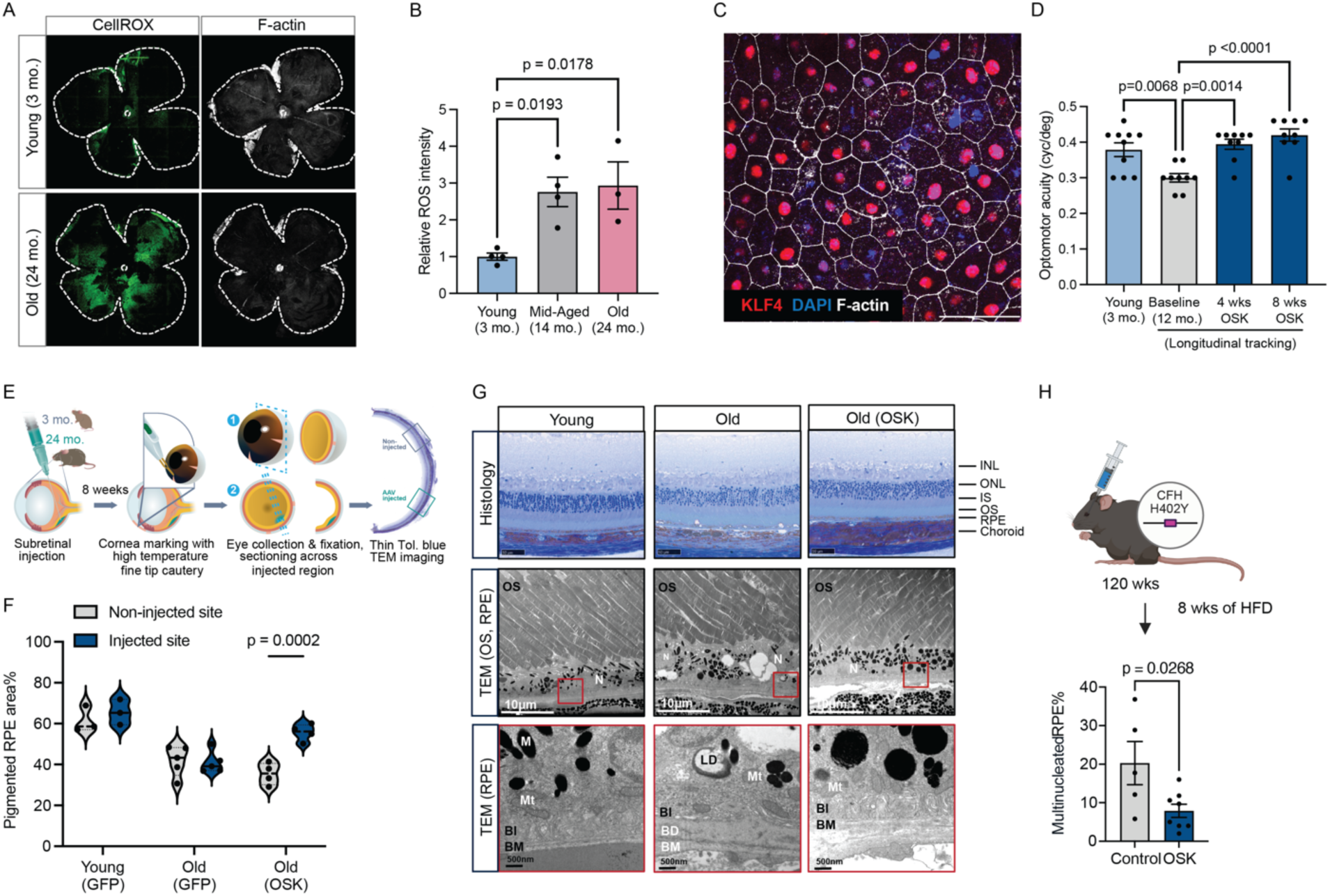
Assessment of OSK-mediated protection against oxidative stress and age-related degeneration in the RPE. (A) Representative RPE flatmounts from young and aged mice stained with CellRox and F-actin. (B) Quantification of relative ROS intensity by immunofluorescence staining in RPE flatmounts from young (3-month-old, 2M2F), middle-aged (14-month-old, 2M2F), and aged (24-month-old, 1M2F) mice (n ≥ 3 eyes per group). One-way ANOVA-Bonferroni. (C) Representative image of AAV-mediated OSK delivery to the subretinal space and transduction of RPE cells. KLF4 (red) serves as a marker of OSK treatment, while F-actin (white) serves as a marker of RPE cells. Scale bar, 50µm. (D) Quantification of visual acuity by optomotor response (OMR) in young (5M5F) and 12-month-old mice at baseline and at 4 and 8 weeks following AAV-OSK injection (5M4F). One-way ANOVA-Bonferroni. (E) Schematic of cornea marking technique to identify the AAV injection site. (F) Quantification of RPE pigmentation in 3- and 24-month-old mice following AAV-GFP or AAV-OSK injection (n ≥ 3 eye). Pigment area was calculated as the percentage of pigmented RPE within the total imaged area using ImageJ (40× magnification, NanoZoomer). Two-way ANOVA-Bonferroni. (G) Toluidine blue–stained retinal sections and transmission electron microscopy (TEM) images showing retinal ultrastructure in young (3mo.), aged, and OSK-treated aged (8 wks post AAV injection at 24mo.) mice. Intracellular lipid droplets (LD), basal deposits (BD) are annotated. Scale bar, 10 μm and 500 nm respectively. Retinal layers: INL, inner nuclear layer; ONL, outer nuclear layer; IS, inner segment; OS, outer segment; RPE, retinal pigment epithelium. Mt, mitochondria; M, melanin granule; N, nuclear; POS, photoreceptor outer segment; BI, basal infoldings; BD, basal deposit; BM, Bruch’s membrane. (H) Quantification of multinucleated RPE cells in 24-month-old CFH-H/H mice administered AAV2-tTA or AAV2-tTA/TRE-OSK 2 weeks ahead of switching to a high-fat, cholesterol-enriched diet for 8 weeks. Multinucleation was defined as ≥3 nuclei per RPE cell (n ≥ 5 eyes). Two-tailed Student’s t-test. Data are mean ± SEM.

To investigate whether partial epigenetic reprogramming can rescue tissues undergoing oxidative damage, we injected our previously described dual AAVs that allows constitutive expression of OSK (AAV2-tTA; TRE-OSK) ^9^ into the subretinal space of 15-month-old mice, allowing OSK expression in the RPE (**Figure 1C**). We chose the AAV2 serotype as it demonstrates high RPE tropism via subretinal injection in a FDA approved gene therapy ^24,25^. We then tracked their visual acuity using the optomotor response (OMR) assay. As seen with the approved RPE gene therapies ^24,25^, visual acuity is primarily driven by high-acuity regions rather than the average function across the entire retina. Therefore, improving even the localized area of the RPE that is transduced by AAV, could be sufficient to enhance OMR scores (**Supplemental Discussion**). Notably, the OSK-treated eyes exhibited significantly improved vision starting at 4 weeks post-injection, and by 8 weeks, visual function was comparable to that of 3-month-old young mice (**Figure 1D**). By contrast, matched control eyes with constitutive expression of GFP (AAV2-tTA; TRE-d2EGFP) remain unchanged from their baseline visual acuity (**Figure S1A**).

This OSK-mediated functional enhancement encouraged us to look further into the potential benefits of OSK treatment on RPE morphology. To distinguish changes in retinal histology and RPE ultrastructure at the AAV injection site, we developed a cornea marking technique to locate the AAV injection site even after enucleation of the eye and during tissue processing **(Figure 1E)**. Aging in the RPE is marked by loss of melanin granules, lipofuscin buildup, basal deposits, Bruch’s membrane thickening, microvilli atrophy, abnormal apical mitochondria with loss of cristae and disorganized basal infoldings^26^ (**Figures 1F-1G**). These changes are also features of AMD donor tissues ^27^, where they are even more pronounced ^28^. The site containing the AAV-OSK treated RPE cells displayed a youthful level of RPE melanin granules (**Figure 1F**), a key indicator of RPE health as adequate melanin helps absorb stray light and reduce photo-oxidative damage. This site also displayed reduced lipid deposits, increased basal infoldings (**Figure 1G**), and mitochondria have restored membrane shape with increased cristae ^26^ (**Figures S1B-S1C**). While aging is a primary risk factor for AMD, genetic risk and lifestyle are also major contributors to the development of AMD. Therefore we tested if AAV-OSK was effective in an AMD mouse model, in which 2-year-old *CFH-H402/H402:Cfh-/-* (*CFH-H/H*) mice carrying the human complement factor H (CFH) Y402H risk allele were placed on a high-fat, cholesterol enriched (HFC) diet ^29^. Notably, AAV2-OSK significantly reduces RPE multinucleation, a measurement of RPE dysmorphia in this mouse model (**Figures 1H** and **S1D**). Collectively, OSK expression in aging RPE demonstrates strong protection against oxidative stress: it restores visual function, improves histological architecture, and protects against aging as well as AMD-associated stressors such as a HFC diet and complement-mediated risk.

Although OSK confers these benefits in RPE without affecting retinal thickness (**Figure S1E**), prolonged expression beyond 8 weeks leads to detectable photoreceptor toxicity measured by electroretinogram (ERG) (**Figure S1F**). Notably, this toxicity was not observed in RGCs even after sustained OSK expression for over a year^14^, likely reflecting differences in cellular context and function ^30^. The enhanced tissue function and delayed toxicity in RPE may result from OSK’s mixed transcriptomic profile: using a transcriptomic aging (tAge) clock ^31^, we observed that OSK expression paradoxically increases tAge in mouse RPE *in vivo* (**Figure S1G**), as it not only ‘rejuvenates’ the cytoskeletal/glycolytic and inflammation gene modules, but also simultaneously activates the interferon signaling and mTOR signaling that have “pro-aging” roles (**Figure S1H**). The combination of strong functional restoration with short-term expression and the toxicity concerns associated with prolonged expression, prompted us to dissect OSK’s mechanism and identify the downstream effectors responsible for its benefits.

### OSK Enhances Mitochondrial Resilience via a Dynamic, Tet2-Independent Mechanism

To identify the molecular mechanism by which OSK protects RPE cells against oxidative stress, we turned to RPE cell culture models. We engineered DOX-inducible lentiviral vectors expressing human OSK and used these to infect the human RPE cell line ARPE-19 (**Figure 2A**), which is highly amenable to lentiviral transduction compared to primary RPE cells and iPSC-RPE cells ^32–34^. Unlike many studies that use ARPE-19 cells in their growth phase ^35–37^, we applied an optimized maturation protocol, including nicotinamide treatment ^38^, and ensured three weeks of maturation, which resulted in RPE cells with morphology that more closely reflects their *in vivo* characteristics (**Figure 2B**). After four days of OSK induction, we exposed the cells to four different oxidative stressors, sodium iodate [NaIO_3_] ^39–41^, oxidized low-density lipoprotein [ox-LDL] ^42^, cigarette smoke extract ^43^, tert-Butyl hydroperoxide [tBH] ^44^, as well as an AMD associated stressor complement^45^, then evaluated cell viability by lactate dehydrogenase (LDH) release and cell replating survival assays. The negative d2EGFP controls offered no protection against any insult and, as a putative positive control, we increased expression of NRF2, a potent activator of a variety of protective pathways against oxidative stress ^46^, which only protected RPE cells against cigarette smoke extract. By contrast, OSK provided significant protection against ox-LDL, tBH, complement, and most robustly, NaIO_3_ (**Figures 2C** and **S2A**). NaIO_3_ is a potent prooxidant that increases mitochondrial ROS (**Figure 2D**), and is widely used in preclinical models of AMD to trigger RPE injury ^39–41^. OSK expression protected mitochondrial respiration from NaIO_3_-induced damage (**Figure 2E**), and functional mitochondria were required for this protective effect (**Figure S2B**). To determine if OSK’s oxidative protection is specific to RPE cells, we expressed OSK via doxycycline in mouse fibroblast cell lines derived from young and aged Rosa26-M2rtTA/Col1a1-tetOP-OKS transgenic mice^9,47^. Consistent with our findings in RPE cells, OSK robustly protected fibroblasts against NaIO_3_-induced oxidative stress (**Figures 2F** and **S2C**). This shows that the oxidative resilience activated by Yamanaka factors is found in at least two cell types and two species.

**Figure 2.**
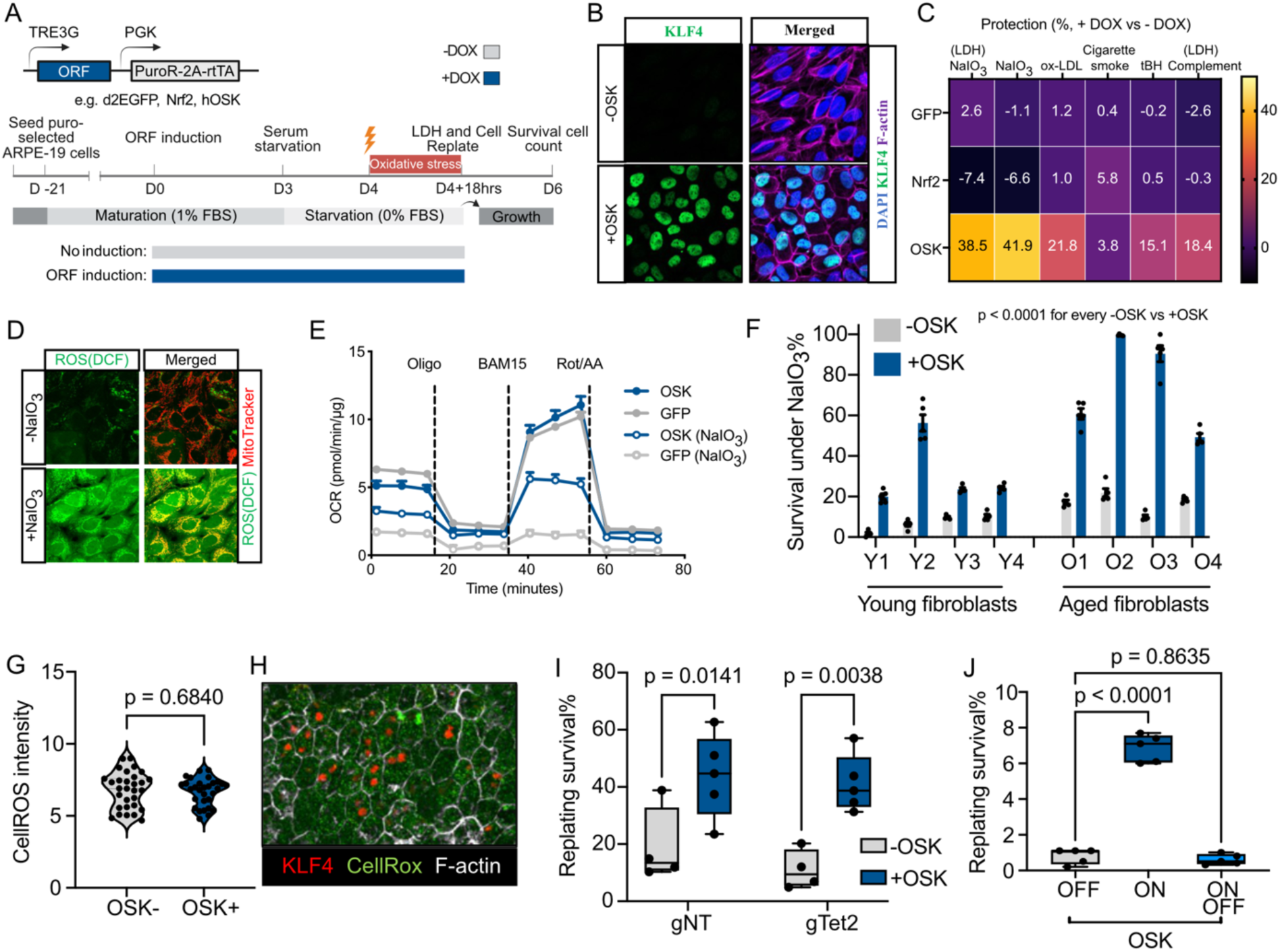
OSK protects mitochondria against oxidative stress through a dynamic, Tet2-independent mechanism. (A) Schematic of the experimental timeline: matured ARPE-19 cells with doxycycline (DOX)-inducible OSK or control ORFs (*e.g.*, d2EGFP, NRF2) were exposed to oxidative stress (5 mM NaIO_3_) and assessed for survival by lactate dehydrogenase (LDH) release and cell replating survival assays. (B) Immunofluorescence images showing morphology of ARPE-19 cells post-maturation, with or without OSK induction, stained for KLF4 (green), F-actin (magenta), and nuclei (DAPI, blue). (C) Heatmap depicting ARPE-19 cell viability under oxidative stress mediated by different stress-inducing conditions. Treatments included addition of either NaIO_3_ for 18 hrs, oxidized low density lipoprotein (ox-LDL) for 24 hrs, cigarette smoke extract for 24 hrs, or tert-Butyl hydroperoxide (tBH) for 4 hrs. Protection was determined by the mean percentage of OSK-expressing cells surviving (+Dox induced OSK) versus the mean percentage of cells surviving without OSK (-Dox) in the replating cell culture survival assay. LDH assays following either NaIO_3_ treatment (LDH NaIO_3_) or complement treatment (LDH complement) represent cytotoxicity levels immediately following treatment (n = 5 in each group). (D) Immunofluorescence images of ROS intensity via DCF staining of ARPE-19 cells in normal culture conditions or following treatment with 5mM NaIO_3_ for 6 hrs. (E) Oxygen consumption rate (OCR) measured via Seahorse Mito Stress Test analysis in OSK-induced and control cells with and without NaIO_3_ treatment (n = 8 in each group). (F) Quantification of mouse fibroblast cell survival with or without OSK induction under oxidative stress conditions (20mM NaIO_3_ treatment for 18 hrs) using the replating cell culture survival assay. Fibroblasts derived from OSK transgenic mice were treated with Doxycycline to induce OSK expression *in vitro* (n = 8 cell lines, Y1-Y4, 3-month-old; O1-O4, 24-month-old). Two-way ANOVA-Bonferroni. (G) Quantification of ROS levels (CellROX) in mouse RPE cells, comparing OSK-expressing and non-expressing RPE cells. Two-tailed Student’s t-test. (H) Representative immunofluorescence image of mouse RPE flatmount from an aged mouse that received a subretinal injection of AAV-OSK 4 weeks prior to recovery and staining for Klf4 (red), ROS (CellROX, green), and F-actin (phalloidin, white). (I) Quantification of ARPE-19 cell survival in the replating cell culture survival assay following Tet2 or control knockout, with (n = 5) or without (n = 4) OSK induction under NaIO_3_ induced oxidative stress conditions. gTet2, CRISPR-Cas9 knockout of Tet2; gNT, non-targeting control. Two-way ANOVA-Bonferroni. (J) Quantification of ARPE-19 cell survival following NaIO_3_ treatment (5mM for 18 hrs). Negative control with OSK off (OFF); positive control with OSK on (ON). RPE cells with OSK on for 4 days followed by withdrawal for 4 days (ON OFF, n = 5). One-way ANOVA-Bonferroni. Data are mean ± SEM.

Next, we investigated whether OSK-induced oxidative resilience follows the canonical pathway. OSK did not directly lower ROS levels in RPE cells (**Figures 2G** and **2H**), suggesting it acts downstream of the ROS response. Our previous work demonstrated the mechanism of OSK-induced rejuvenation of retinal ganglion cells was dependent on Tet2 mediated CpG demethylation ^9^, and supported by another study inhibiting DNMT3a methylation to achieve similar effect ^48^; we therefore tested whether the oxidative protection in OSK-treated RPE cells is mediated through this same Tet2 dependent pathway. Surprisingly, neither CRISPR-Cas9 knockout of Tet2 (**Figures 2I** and **S2D-S2E**) nor widely used Tet2 knockdown by shRNA (**Figures S2F-S2G)** blocked OSK-mediated protection in NaIO_3_-treated RPE cells. Consistent with the absence of stable DNA methylation-associated memory, the oxidative protective effect requires continuous OSK expression: transient OSK expression followed by withdrawal of OSK failed to maintain oxidative protection in NaIO_3_-treated RPE cells (**Figures 2J** and **S2H-S2I**). Collectively, these findings demonstrate that sustained OSK expression triggers a non-canonical, Tet2-*independent* resilience program that safeguards cellular function against oxidative insults.

### GSTA4 Is a Necessary and Sufficient Downstream Effector of OSK-Mediated Oxidative Protection

The above data demonstrated that OSK confers anti-oxidative benefits across three distinct model systems: aged mouse RPE cells under age-elevated oxidative stress, and human RPE cells and mouse fibroblasts exposed to oxidative stressors *in vitro*. To uncover the common molecular mechanisms underlying these effects, we generated three comprehensive transcriptomic datasets: (1) human ARPE-19 cells overexpressing OSK or GFP for four days; (2) RPE cells from aged mice expressing AAV-OSK or AAV-tTA for five weeks; and (3) young and aged mouse fibroblast lines with or without OSK expression for six days (**Figure 3A**). To identify potential downstream effectors, we first pinpointed differentially expressed genes (DEGs) shared across all three models (**Figure 3B**) and assessed whether their expression correlated with survival rates against NaIO_3_-induced oxidative stress (**Figure 3C**). This analysis led us to nominate six candidate effectors: Gsta4, Aldh3a1, Pipox, Ccdc3, Nrarp, Phyhip (**Figures 3C and S3A**), based on their strong induction by OSK across all three systems and tight correlation with oxidative protection. We then cloned each of these candidates, along with the poorly correlated Carns1 as a negative control, into doxycycline-inducible vectors to create stable ARPE-19 lines. After three weeks of maturation, we induced overexpression for four days and performed an arrayed screen to evaluate their ability to protect against NaIO_3_-induced oxidative stress (**Figures 2A** and **3D**).

**Figure 3.**
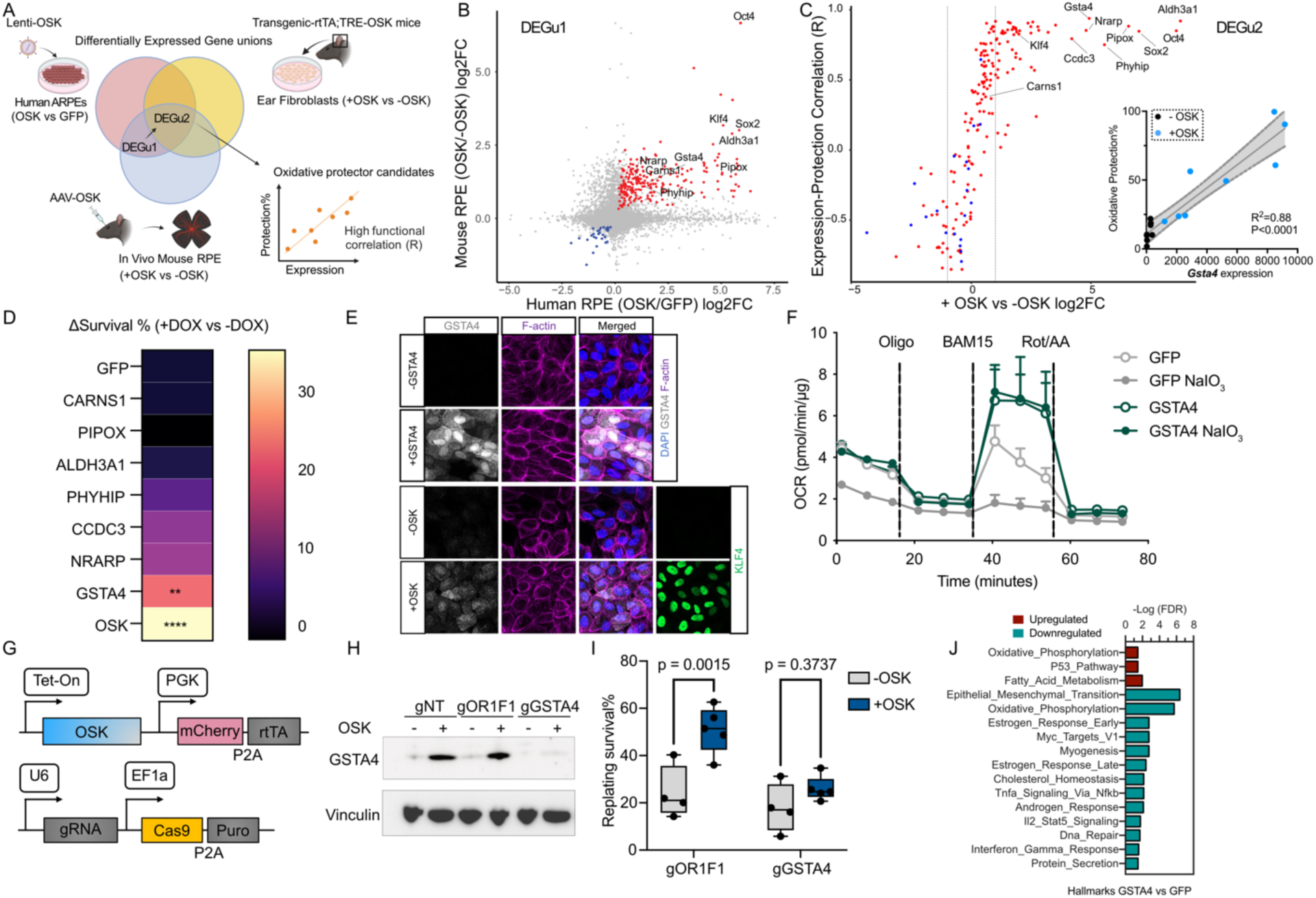
Identification of GSTA4 as a necessary and sufficient OSK-downstream effector responsible for oxidative resilience. (A) Overview of transcriptomic datasets used to identify OSK-regulated gene unions in human ARPE-19 cells (OSK vs GFP, 4 days), aged mouse RPE (AAV-OSK vs AAV-tTA, 5 weeks post injection in 20-month-old mice), and mouse fibroblasts (young and aged, 8 cell lines, ±OSK for 6 days). (B) Volcano plots showing differentially expressed gene union 1 (DEGu1) induced by OSK expression in aged mouse RPE cells and human ARPE-19 cell transcriptomic datasets. (C) Scatter plot of the shared DEGs across two RPE systems (as in Figure 3B), mapped onto the mouse fibroblast data. Each gene is positioned by its OSK-induced log₂ fold change (x-axis) in fibroblast and its Pearson correlation (y-axis) between gene expression and NaIO_3_ protection (as in Figure 2F). Gsta4 correlation figure is exemplified at the lower right corner, with –OSK (black) and +OSK (blue) points illustrating its expression levels and oxidative protections. Shaded regions indicate 95% confidence intervals around the linear regression line. Correlation figures of additional candidates are displayed in **Figure S3A**. (D) Heatmap depicting ARPE-19 cells mean viability following individual overexpression of selected candidate ORFs in the context of oxidative stress. (E) Immunofluorescence images of GSTA4 protein in ARPE-19 cells. Staining is shown for control cells (-OSK), OSK-induced cells (+OSK), and cells with direct GSTA4 overexpression, stained for GSTA4 (white), F-actin (magenta), KLF4(green) and nuclei (DAPI, blue). (F) Oxygen consumption rate (OCR) measured via Seahorse analysis in GSTA4-induced and control cells (GFP) before and after NaIO_3_ treatment. (G) Schematic of the inducible OSK overexpression and CRISPR-Cas9 system used to disrupt GSTA4 expression. (H) Immunoblot analysis of GSTA4 protein level under OSK induction in CRISPR-mediated knockout ARPE-19 cells. (I) Quantification of ARPE-19 cells survival following CRISPR-Cas9 knockout of GSTA4 compared to controls (gOR1F1), with (n = 5) or without (n = 4) OSK induction under oxidative stress conditions. Two-way ANOVA-Bonferroni. (J) Gene set enrichment analysis (GSEA) of transcriptomic data following OSK expression, highlighting oxidative phosphorylation, fatty acid metabolism, and epithelial–mesenchymal transition–related gene signatures. All data are presented as mean ± SEM.

Among the candidate genes, GSTA4 emerged as the *sole* factor that was sufficient to provide significant oxidative protection to RPE cells (**Figures 3D,3E** and **S3B**) and successfully recapitulated OSK’s ability to protect mitochondrial function under NaIO_3_-induced oxidative challenge (**Figure 3F)**. Immunostaining and Western blot analysis confirm OSK up-regulating GSTA4 at the protein level, and importantly, CRISPR-Cas9-mediated knockout of GSTA4 abolished OSK’s protective effects against NaIO_3_-induced oxidative stress (**Figures 3G-I** and **Figure S3C**), while knockout of a non-expressed control gene OR1F1 does not (**Figure 3I**). In aging and disease, RPE cells down-regulate oxidative phosphorylation and up-regulate epithelial-to-mesenchymal transition (EMT) pathways ^49,50^. At the transcriptomic level, overexpression of either OSK or GSTA4 alone in ARPE-19 cells reverses these effects; both upregulating oxidative phosphorylation and down-regulating EMT (**Figures 3J** and **S3D**). Together, we identify GSTA4 as a necessary and sufficient downstream effector of OSK, mediating its oxidative resilience and mitochondrial protective effects.

### OSK Reactivates a Developmentally Regulated GSTA4 Stress Resistance Pathway

To understand how the three Yamanaka factors (OSK) regulate GSTA4, we first examined whether any of the individual factors could induce its expression. Interestingly, each individual factor produced a modest rise in GSTA4, an effect unique among the candidates, while their combined expression markedly amplified GSTA4 induction (**Figure 4A**). Why do all three stem-cell factors converge on GSTA4? We suspect this redundant activation safeguards stem cells by providing critical oxidative resilience. To examine its functional relevance, we revisited our published CRISPRi screen dataset in human stem cells ^51^; analysis of this unbiased dataset showed that GSTA4 knockdown indeed significantly diminishes stem-cell fitness and growth, underscoring its essential oxidative protective role (**Figure 4B**).

**Figure 4.**
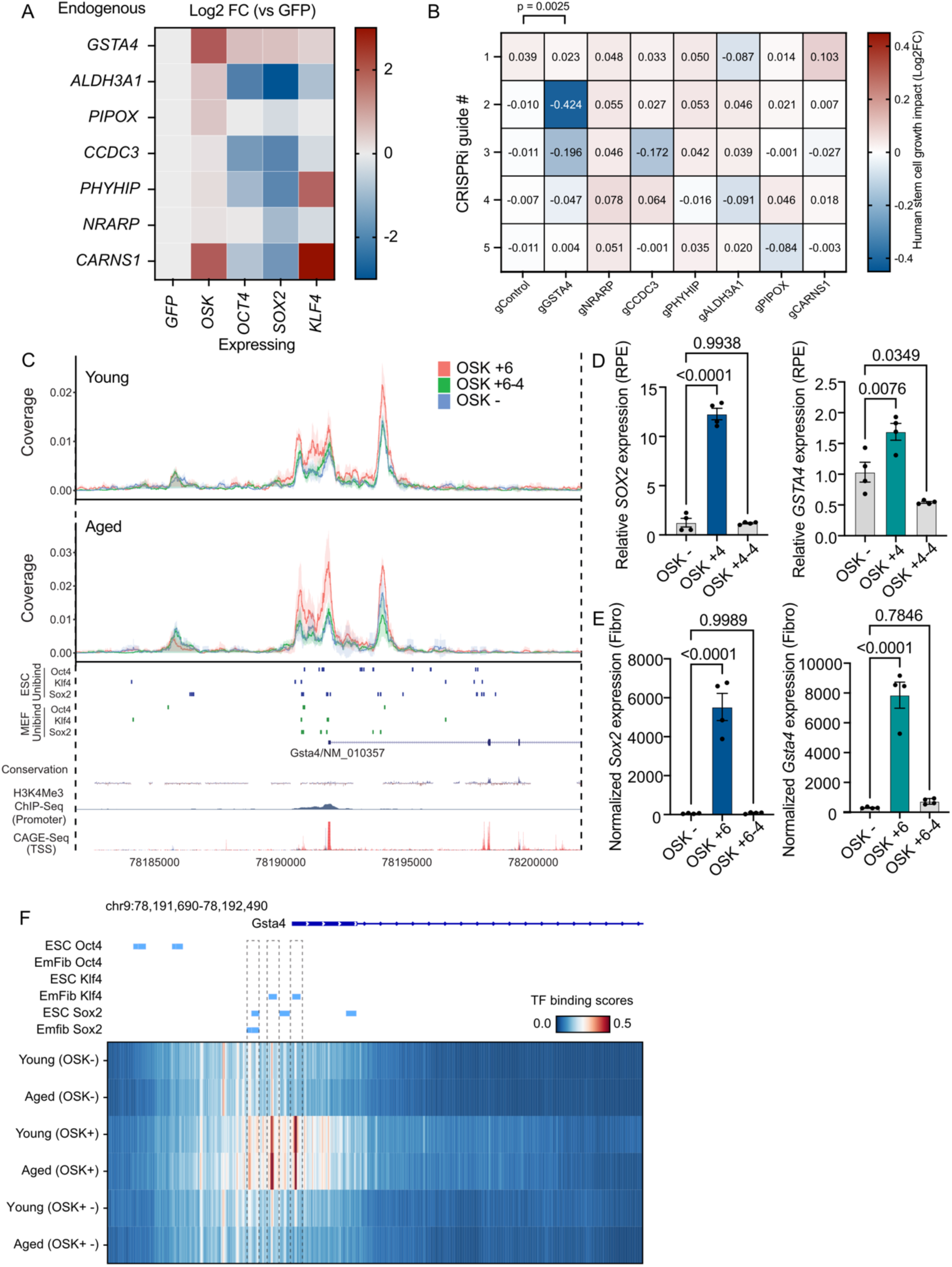
GSTA4 is a direct and dynamic target of OSK factors and is essential for stem cell function. (A) Heatmap showing induction of candidate oxidative effectors upon ectopic expression of Oct4, Sox2, Klf4 or OSK in ARPE-19 cells. (B) Stem cell growth impact of sgRNAs targeting candidate genes using CRISPR interference (CRISPRi)-mediated knockdown in human embryonic stem cells, represented as the average Log₂ fold change (Day 10/Day 0) in WTC11 and H1 hESC lines. (C) Chromatin accessibility and UniBind-based transcription factor binding profiles for Oct4, Sox2, and Klf4 at the Gsta4 locus under continuous or transient OSK expression, with H3K4me3 ChIP-seq serving as a promoter mark and CAGE-seq indicating the transcription start site. Unibind ChIP-validated TF binding sites obtained from mouse embryonic stem cells (ESCs) and mouse embryonic fibroblasts (EmFib). +6, OSK induced with Doxycycline for 6 days; +6-4, OSK induced with doxycycline for 4 days, then withdrawn for 4 days; -, no OSK induction. (D and E) mRNA transcript levels of GSTA4 and Gsta4 under continuous or transient OSK expression in human RPE (D) and in mouse fibroblasts (E). Two-way ANOVA-Bonferroni (n = 4 per group). +4, OSK induced with doxycycline for 4 days; +4-4, OSK induced with doxycycline for 4 days, then withdrawn for 4 days; -, no OSK induction. (F) TF binding score figure: Top tracks: Unibind ChIP-validated TF binding sites obtained from mouse embryonic stem cells (ESCs) and mouse embryonic fibroblasts (MEFs). Overlapping sites are merged. Bottom heatmap: seq2PRINT-inferred TF binding scores within the Gsta4 promoter region at chr9:78191690-78192490. Columns represent single base pair positions, and rows represent individual samples. All data are presented as mean ± SEM.

Considering the OSK-induced protection from oxidative stress does not sustain (**Figure 2J**), we suspect that OSK-mediated upregulation of GSTA4 is acute but also dynamic. To examine this, we performed ATAC-seq in cells with either sustained or transient OSK expression. As expected, when OSK was expressed, chromatin accessibility at the GSTA4 promoter increased, and it returned to a closed state once OSK was removed (**Figure 4C**). This dynamically regulated chromatin accessibility parallels GSTA4 RNA expression levels in both human RPE cells and mouse fibroblasts, where its RNA expression significantly declines upon OSK withdrawal (**Figures 4D**, **4E**). Notably, the newly accessible chromatin region at the GSTA4 promoter harbors binding motifs for all three factors Oct4, Sox2 and Klf4 (**Figure 4C**). To map factor binding at a high resolution, we applied a novel deep-learning–based seq2PRINT method ^52^ that allows precise inference of transcription factor binding. seq2PRINT inferred significantly increased Klf4 and Sox2 binding at the proximal Gsta4 promoter region during OSK overexpression. Importantly, upon OSK withdrawal, such occupancy returns to baseline (**Figure 4F**). This rapid remodeling at the Gsta4 promoter parallels the dynamic GSTA4 induction, and aligns with the Tet2-independent, memory-free nature of the OSK-induced oxidative protection.

The above observations raise the question of why the activation of GSTA4 occurs in a rapid and reversible manner. We propose that it might be important for GSTA4 to be down-regulated as stem cells exit pluripotency, permitting the ROS surge that drives mesendoderm differentiation — an idea supported by evidence that elevated ROS is essential for this lineage commitment ^53^. To explore this, we performed more detailed analysis of single-cell datasets from embryonic stem cells ^54^ (**Figure S4A** and **S4B**). The analysis revealed high GSTA4, Oct4, Sox2 expression post implantation, followed by a sharp decline during mesendoderm commitment (**Figures S4C** and **S4D**). It also coincides with the activation of EMT genes that are known drivers of mesoderm formation (**Figure S4E**). This observation suggests a potential evolutionary advantage to the dynamic regulation of GSTA4: a swift downregulation of GSTA4 following OSK withdrawal may permit a ROS surge that facilitates mesendoderm formation, in line with evidence that ROS scavengers inhibit this process ^53^. Taken together, our analysis reveals dynamic GSTA4 regulation by OSK, and suggests that this protective mechanism may be programmed to turn off during development-unless reactivated through ectopic OSK expression.

### AAV-Mediated GSTA4 Expression Improves Vision and Reduces 4-HNE in Aging RPE

Building on the *in vitro* benefits, we next asked whether GSTA4 alone could reproduce OSK’s *in vivo* effects (**Figure 1D**). We packaged GSTA4 in an AAV2 vector and, after recording baseline OMR in 15-month-old mice, delivered subretinal AAV-GSTA4 to one cohort and AAV-GFP to controls (**Figure 5A**). Notably, the GSTA4-treated eyes showed significantly improved visual function beginning at 6 weeks post-AAV injection with progressive improvements by 8 weeks (**Figure 5B** and **Supplemental Videos**). No toxicity or detriment to retinal thickness was observed following 8 weeks of GSTA4 overexpression (GSTA4-OE) in RPE cells (**Figures S5A-S5B**), nor was any photoreceptor-layer toxicity detected by full-field ERG (**Figure S5C**), indicating a safer profile than prolonged OSK expression (**Figure S1F**).

**Figure 5.**
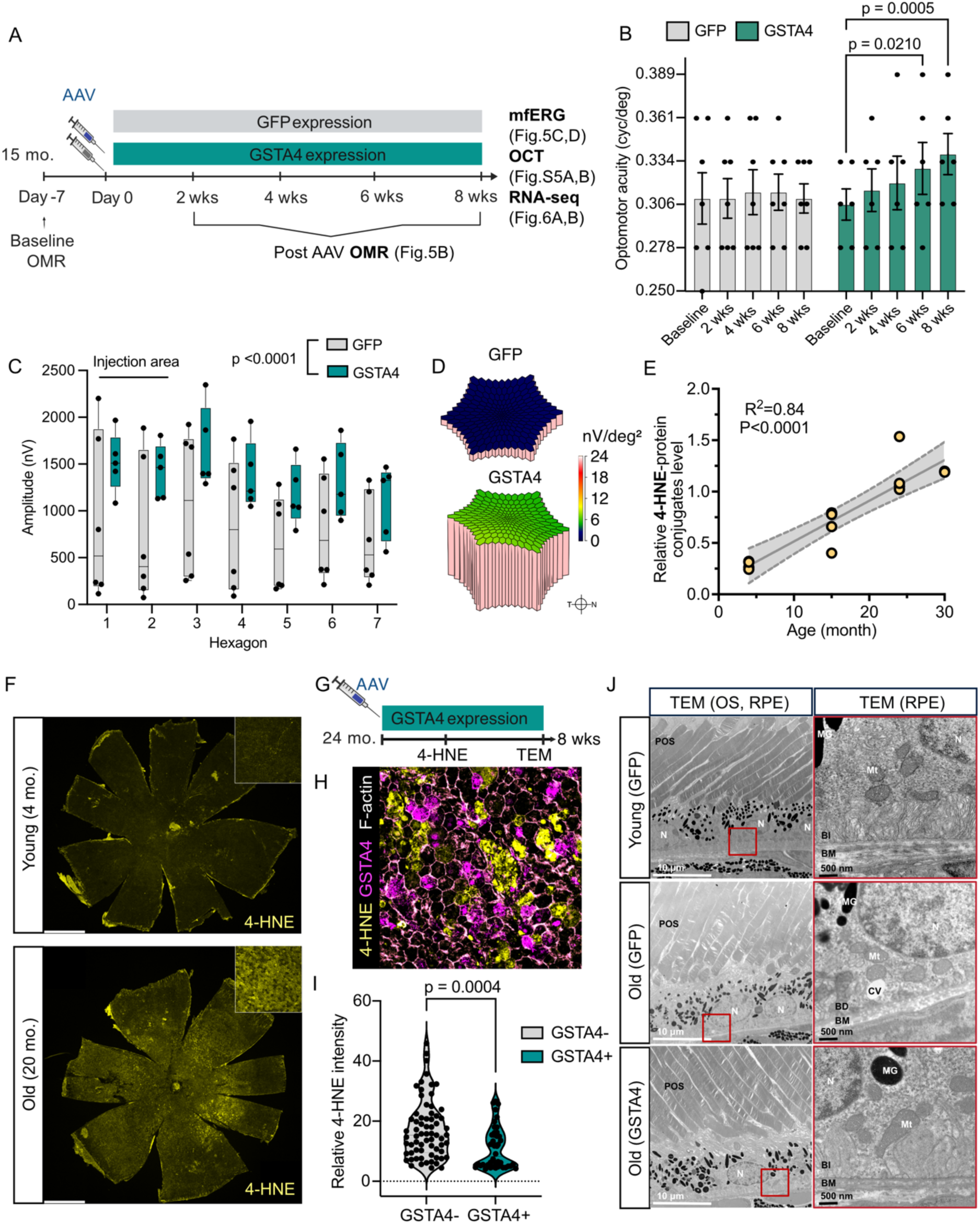
AAV-GSTA4 targets age-related 4-HNE accumulation and enhances visual function and retinal electrophysiology in aged mice. (A) Schematic of the experimental timeline for subretinal AAV injection and subsequent measurements, including optomotor response (OMR), optical coherence tomography (OCT), and multifocal electroretinography (mfERG) in 15-month-old mice. (B) Quantification of visual acuity by OMR at baseline and at indicated time points following AAV injection (GFP group, n = 7, 4M3F; GSTA4 group, n = 6, 3M3F). The visual acuity examiner was masked to the mouse treatment. Two-way ANOVA-Bonferroni. Representative before and after optomotor recordings are provided in **Supplemental Videos**. (C) Multifocal electroretinogram (mfERG) responses recorded in AAV-treated eyes at 8-weeks post-AAV treatment (n = 6 in GFP group, n = 5 in GSTA4 group with one failed to sleep). mfERG testing uses a stimulus pattern across the retina made up of multiple hexagons, each with a unique stimulus, allowing the system to determine the electrical response from specific areas of the retina. Two-way ANOVA-Bonferroni. (D) The three-dimensional plots depicting representative responses of GFP and GSTA4 groups in mfERG, plotted by RMS Density x nV/deg² (nV per square degree). A higher score indicates better retinal function. (E) Scatter plots showing the correlation between chronological age and 4-HNE–protein conjugates levels in the retina–RPE, normalized to actin. Shaded regions indicate 95% confidence intervals around the linear regression line. (F) Representative immunofluorescence image of RPE flatmounts from young (4-month-old) and old (20-month-old) mice stained for 4-HNE-protein conjugates. Corresponding F-actin staining is shown in **Figure S5E**). Scale bar, 1mm. (G) Schematic of the experimental timeline for accessing GSTA-OE in alleviating 4-HNE and ultrastructure dysfunction in aged retinas (24 mo.). (H) Representative immunofluorescence image of RPE from 24-month-old mice, three weeks after subretinal AAV-GSTA4 delivery, stained for 4-HNE (yellow), GSTA4 (purple) and F-actin (phalloidin, white). (I) Quantification of relative 4-HNE fluorescence intensity in GSTA4⁺ and GSTA4⁻ cells across eight sampled fields per group, from 24-month-old mice, three weeks after subretinal AAV-GSTA4 delivery. Two-tailed Student’s t-test. (J) Transmission electron microscopy (TEM) images showing retinal ultrastructure in GFP-treated young (5 mo.), GFP-treated aged, and GSTA4-treated aged mice (26 mo., 8 weeks post AAV-GSTA4). Procedure follows demonstration in Figure 1E. Basal deposits (BD), and cytoplasmic vacuolization (CV) are annotated. Scale bar, 10 μm and 500 nm respectively. Mt, mitochondria; M, melanin granule; N, nuclear; POS, photoreceptor outer segment; BI, basal infolding; BD, basal deposit; BM, Bruch’s membrane. All data are presented as mean ± SEM.

To better assess localized functional gains, we applied a light adapted multifocal electroretinogram (mfERG) system, which stimulates many retinal areas simultaneously and records localized responses, overcoming full-field ERG’s limitation of capturing only a summed signal from the entire retina (**Supplemental Discussion**). The recordings indicated enhanced retinal electro-function to light in GSTA4 overexpressing eyes, with higher electrical responses to light stimuli near the AAV-GSTA4 injection site (**Figures 5C** and **5D**).

GSTA4 is a member of the glutathione S-transferase family that plays a crucial role in detoxifying 4-hydroxynonenal (4-HNE), a harmful byproduct of lipid peroxidation ^55^. Because oxidative stress rises with RPE aging (**Figures 1A** and **1B**) and lipid peroxidation is observed in RPE dysfunction^28^, we assessed 4-HNE levels in young and old eye samples. We observed a progressive, age-correlated increase in 4-HNE within the retina-RPE (**Figures 5E** and **S5D**), with significantly higher 4-HNE accumulation in aged RPE flatmounts compared to young (**Figures 5F** and **S5E**), supporting 4-HNE as a potential age-related biomarker for RPE.

To test if GSTA4-OE could benefit at even older age, we subretinally injected AAV2-GSTA4 into eyes of 24-month-old mice. Through flatmount staining, we observed that aged RPE cells overexpressing GSTA4, compared to those not, exhibited significantly reduced levels of 4-HNE (**Figures 5G-5I**). This aligns with GSTA4’s role in detoxifying 4-HNE by conjugating it with glutathione to form the more soluble, less toxic GSH-HNE that can then be exported from the cell. At ultrastructural morphology, similar to AAV-OSK (**Figure 1G**), AAV-GSTA4-treated eyes showed less evident lipoprotein-rich basal deposits (BD) and cytoplasmic vacuolization (CV) compared to control aging eyes (**Figure 5J**), while the number of cristae in RPE mitochondria was restored to youthful level (**Figures S5F-S5G**), demonstrating an improvement in RPE health. Together, we identified 4-HNE as an aging biomarker of RPE cells, and that enhancing GSTA4 expression bolsters the cellular defenses in aged RPE against toxic 4-HNE, reduces oxidative damage, and restores RPE function and vision, supporting a causal role of 4-HNE in age-related RPE degeneration and GSTA4-OE as a promising therapeutic intervention.

### Reactivation of GSTA4 in Aged RPE Recapitulates the Molecular Signatures of Longevity Interventions

Since we observed that GSTA4 treatment improves visual function in aged mice, we next asked whether it likewise rejuvenates the transcriptome of aged RPE cells. Following 8 weeks of GSTA4-OE, we collected RPE cells at 17-month-old and performed RNA-seq, along with young and aged controls. Using a previously developed multi-tissue transcriptomic aging clock ^31^, we detected a pronounced aging signature in the young and aged control RPEs (**Figure 6A**), marked by upregulation of inflammatory and complement pathway genes (**Figure 6B**). Remarkably, GSTA4-OE reduced the transcriptomic age of aged RPE cells by 8.5 months – approximately 50% of their chronological age – and induced a gene expression profile associated with reduced expected mortality (**Figures 6A** and **S6A)**. GSTA4 treatment shifted the transcriptome toward a state more closely resembling young cells, particularly in the cytoskeletal, glycolytic, and inflammatory gene modules (**Figure S6B)**, similar to OSK. Importantly, this occurred without activation of interferon or mTOR signaling, which are induced by OSK (**Figure S1H**). As a result, GSTA4-OE reduced expression of inflammatory and complement pathway genes (**Figure 6B**) while upregulating genes involved in mitochondrial function and fatty acid metabolism (**Figure S6C**).

**Figure 6.**
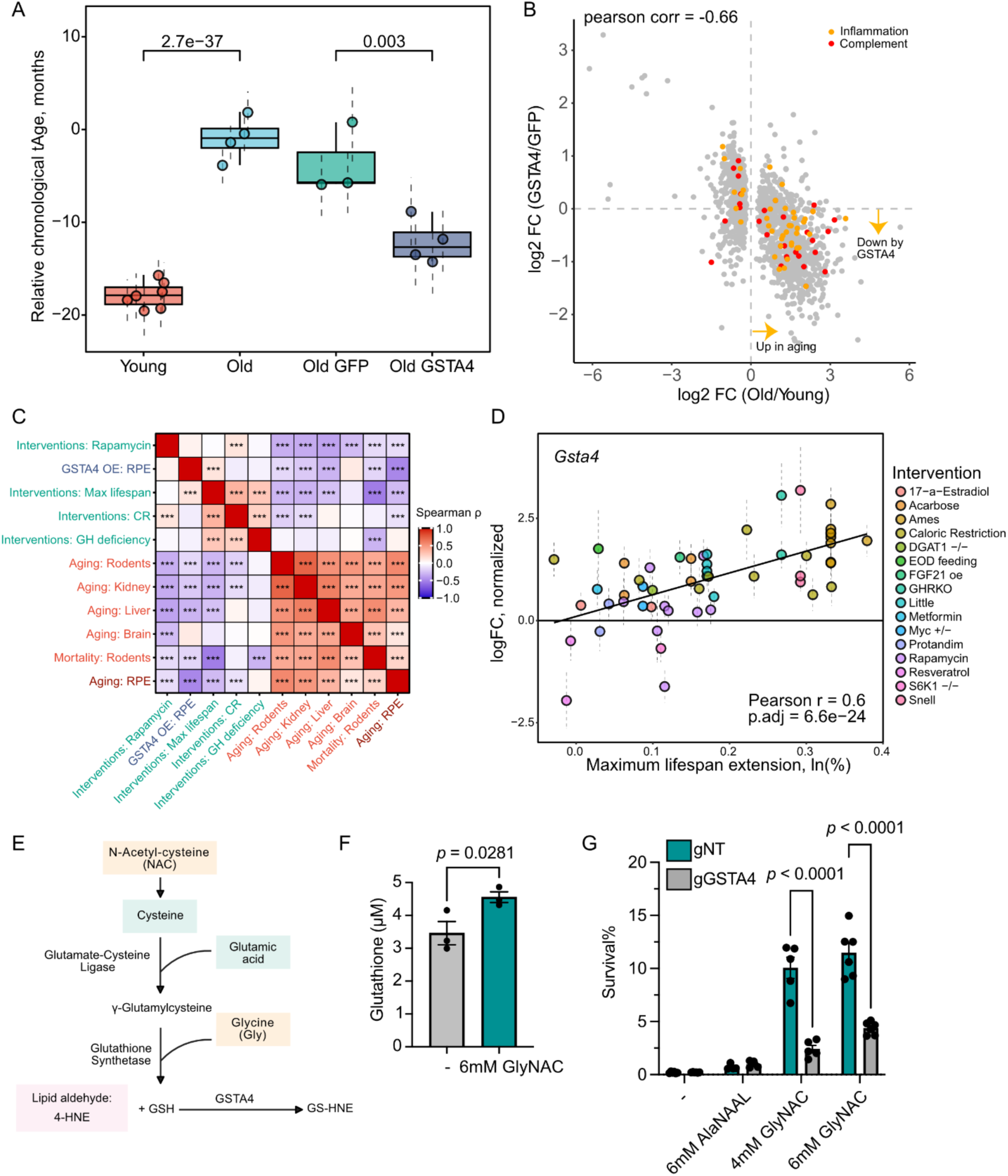
GSTA4 overexpression in RPE restores youthful gene expression, links to lifespan-extending treatments, and mediates GlyNAC’s protective effect. (A) Transcriptomic age (tAge) of young (4 mo.) and old (17 mo.) mouse RPE cells, untreated and with AAV-GSTA4 treatment, as predicted by multi-tissue transcriptomic clock of chronological age. (B) Log2 fold change of gene expression in RPE aging and after AAV treatment. (C) Spearman correlation heatmap comparing gene expression signatures of GSTA4 AAV, mortality, aging (in RPE, brain, liver, kidney, and across tissues) and established lifespan-extending interventions (growth hormone deficient models, caloric restriction, rapamycin, and signature of maximum lifespan extended by various interventions). Each signature corresponds to a vector of normalized enrichment scores (NES) derived from gene set enrichment analysis (GSEA) using the HALLMARK, KEGG, and REACTOME ontologies. *** p.adj < 0.001; ** p.adj < 0.01; * p.adj < 0.05. (D) Positive associations between hepatic expression change of Gsta4 and the effect of longevity interventions on mouse expected maximum lifespan. Each dot corresponds to a single dataset and represents the mean logFC of the gene in response to a particular treatment. Error bars are standard errors (SE). (E) Schematic showing GSTA4 as the key mediator of GlyNAC’s cytoprotective effect. (F) Quantification of glutathione levels in ARPE-19 cells following supplementation with 6mM GlyNAC (Glycine and N-Acetyl-L-cysteine, n = 3 per group). Two-tailed Student’s t-test. (G) Quantification of survival following NaIO_3_ challenge in ARPE-19 cells with CRISPR-Cas9 knockout of GSTA4 (gGSTA4) compared to non-targeting controls (gNT), under no treatment (-) or treatment with either 4 mM GlyNAC (Glycine and N-acetylcysteine), 6 mM GlyNAC or 6 mM AlaNAAL (L-Alanine and N-Acetyl-L-alanine, n = 5 per group). Two-way ANOVA-Bonferroni. All data are presented as mean ± SEM.

To assess the broader relevance of GSTA4-induced transcriptomic rejuvenation, we compared gene expression changes in GSTA4-treated aged RPE cells with transcriptomic signatures of aging across multiple tissues and of established lifespan-extending interventions^56^. The resulting correlation heatmap shows that RPE aging strongly resembles transcriptomic changes in other aged tissues (*e.g.* brain, liver, kidney) and those associated with increased mortality in rodents. Notably, AAV-GSTA4-treated RPE exhibits strong negative correlation with aging signatures, and positive correlation with gene expression profiles associated with extended maximum lifespan induced by multiple longevity interventions, such as caloric restriction (CR), genetic models of growth hormone deficiency, and rapamycin (**Figure 6C**).

Using the previously developed mSALT tool ^56^, which integrates gene expression responses to various lifespan-expanding interventions in mouse liver, we found GSTA4 to be significantly associated with murine maximum lifespan (**Figure 6D**) and upregulated across eight individual longevity interventions, including CR and GHRKO – two of the most robust models of lifespan extension – as well as several pharmacological pro-longevity treatments, including metformin, acarbose, and 17α-estradiol (**Figure S6D**). When ranked by association between the expression level and the magnitude of lifespan extension, GSTA4 ranked as one of the strongest lifespan-associated genes, placing among the top 0.71% genes by statistical significance and among the top 0.45% by normalized slope of association with maximum lifespan across interventions (**Figure S6E**). Because RPE aging parallels that of other tissues (**Figure 6C**) and hepatic GSTA4 induction is a common feature of multiple lifespan-extending interventions, our findings raise the notion that reactivating GSTA4 in RPE engages conserved systemic anti-aging pathways with the potential to safely promote longevity beyond the eye.

### Glutathione Precursors Restores Oxidative Resilience via a GSTA4-Dependent Pathway

Given that GSTA4 detoxifies reactive aldehydes by conjugating them to glutathione, we asked whether supplementing its substrate could likewise bolster oxidative resilience. Aging is accompanied by glutathione depletion in human and mouse ^57^, yet direct supplementation of glutathione often perturbs cellular redox homeostasis and can be deleterious. By contrast, GlyNAC, a combination of two glutathione precursors glycine and N-acetylcysteine, supports endogenous synthesis (**Figure 6E)** and has been shown to extend lifespan in 15-month-old mice ^58^, and improve metabolic and functional health in elderly humans ^59^. We hypothesize that supplementing cells with GlyNAC could accelerate the GSTA4-mediated detoxification process, effectively mimicking the effects of GSTA4 upregulation. We confirmed that six-hour GlyNAC treatment significantly elevated intracellular glutathione in RPE cells *in vitro* (**Figure 6F)**. Indeed, GlyNAC co-treatment conferred robust protection against NaIO_3_-induced oxidative challenge, an effect that was abolished in GSTA4-knockout cells, demonstrating a GSTA4-dependent mechanism (**Figures 6G** and **S6F**). GlyNAC did not induce cell proliferation when unchallenged, so the effect cannot be explained by proliferation (**Figure S6G**). To rule out simple extracellular quenching of NaIO_3_, we tested structurally analogous compounds, alanine and N-acetyl-L-alanine, which are not substrates for glutathione biosynthesis and these did not offer significant protection against NaIO_3_ (**Figure 6G)**. Together, these results identify GSTA4 as the key mediator of GlyNAC’s cytoprotective effect *in vitro*, and suggest that supplementing precursors of glutathione cofactor for GSTA4 may restore oxidative resilience in RPE, encouraging future investigations to treat AMD.

## DISCUSSION

Over the past decade, partial epigenetic reprogramming has captured the attention of the aging field by demonstrating that transient expression of some or all of the Yamanaka factors can reset cellular age, restore function across diverse tissues and extend lifespan ^8–13^. However, the multifaceted nature of the response to overexpression of Yamanaka factors, which impacts the expression of thousands of genes and imparts permanent changes to the epigenetic state of a cell, poses a challenge to efforts to understand the rejuvenation process or to use partial reprogramming for broad clinical benefit, particularly in the tissues and cell types where toxicity has been reported with prolonged OSKM expression^12,21^.

Ideally, for each tissue application one would identify and utilize the specific OSK(M) downstream effectors—functional units that directly carry out the beneficial biological processes. This has been a challenging task due to the lack of a readily screenable aging phenotype and the limited passage capacity of primary aged cells, restricting unbiased genetic screens, except in the case of aged stem cells that can be expanded ^60^. Focusing on aging in RPE cells, key retinal cells exposed to high oxidative stress whose decline contributes to AMD, we set out to address this question by identifying a screenable phenotype and narrowing the scope of the screen using integrative datasets and functional correlation-based candidate selection.

Here, we first characterize a novel OSK-induced trait: enhanced oxidative resilience in RPE both in vitro and in vivo, mediated by a non-canonical, Tet2-independent mechanism that does not leave a lasting memory (**Figure 7A**). RNA-seq analysis of three relevant cellular systems that displays oxidative resilience upon OSK induction — human ARPE-19, primary mouse RPE and fibroblasts — help reveal a core set of shared differentially expressed genes. By correlating each gene’s induction magnitude with the degree of oxidative resilience, then performing an overexpression screen, we identified GSTA4 as a key effector sufficient to largely recapitulate OSK’s oxidative defense. Using CRISPR-based gene knockouts, we further demonstrated that GSTA4 is necessary for OSK-mediated protection (**Figure 7B**). Most notably, GSTA4-OE in aged RPE rejuvenates transcriptomic aging profile and improves visual functions—without the long-term toxicity seen with OSK (**Figure 7C**). To our knowledge, GSTA4 is the first identified downstream effector of partial reprogramming. Further exploration of OSK-regulated genes linked to its benefits in other cellular contexts has a high chance to uncover additional effectors.

**Figure 7.**
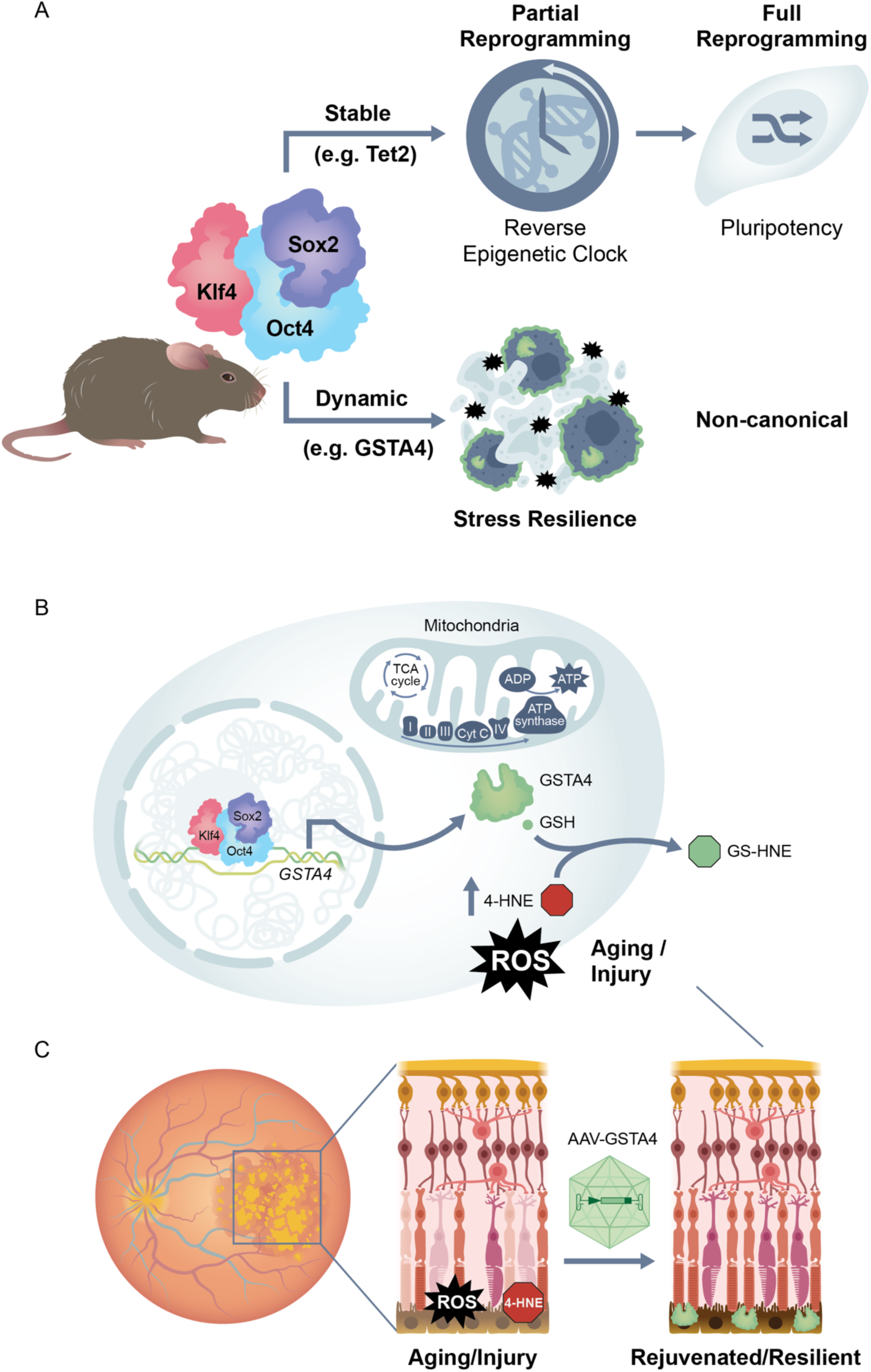
The OSK-GSTA4 pathway exemplifies a dynamic, Tet2-independent stress-resilience axis that can be harnessed to treat age-related diseases. (A) Schematic highlighting the dual outcomes of OSK-mediated reprogramming. Partial reprogramming with OSK is traditionally viewed as resetting the epigenetic clock via modifiers like Tet2. We now show that in RPE cells and fibroblasts, OSK also dynamically up-regulate stress-resilience factors, most notably the detoxifying enzyme GSTA4, through a Tet2-independent manner, thereby strengthening cellular defenses. (B) Schematic of OSK action in RPE cells. Transient expression of OSK engages a Tet2-independent pathway that directly up-regulates the detoxifying enzyme GSTA4 without inducing full epigenetic reprogramming. GSTA4, together with its co-substrate glutathione (GSH), catalyzes conjugation and export of the lipid peroxidation product 4-HNE, thereby preserving mitochondrial electron transport chain function under oxidative stress. (C) In aged or injured retinas, elevated ROS and 4-HNE accumulation in the RPE (middle) disrupt photoreceptor alignment and visual function. AAV-mediated delivery of GSTA4 restores RPE mitochondrial health, clears 4-HNE, reverses age-associated transcriptional drift and rescues retinal structure and vision (right), offering a more precise and safer strategy to treat RPE aging and potentially AMD.

GSTA4 plays a crucial role in detoxifying 4-HNE, the toxic byproduct of lipid peroxidation which has been implicated in a wide range of human diseases including cancer, diabetes, cardiovascular and inflammatory complications ^61^. Here we show that aged RPE cells accumulate ROS and that 4-HNE rises progressively with age (**Figures 5E** and **5F**). Together with our finding of the functional benefits from GSTA4-OE, these observations provide additional support for the proposal that oxidative stress damage acts as a primary driver of RPE dysfunction during aging. Similarly, 4-HNE accumulation is characteristic of other neuro-degenerative disorders: its levels are elevated in tissues and fluids from patients with Alzheimer’s disease (AD) and ALS, and 4-HNE adducts localize to key pathological structures—such as neurofibrillary tangles and plaques in AD, residual motor neurons in ALS, and Lewy bodies in Parkinson’s disease and diffuse Lewy body disease ^62^. Therefore, the consistent up-regulation of GSTA4 across multiple lifespan-extending interventions suggests potential benefits for brain aging and supports its favorable safety profile as a therapeutic candidate for broader application in age-related disorders marked by 4-HNE accumulation.

A recent study reported broad mesenchymal drift across aging and multiple diseases, and further showed that partial epigenetic reprogramming can reverse this process ^63^. Our findings in RPE aging resonate with this work, and further suggesting that oxidative stress and the accumulation of 4-HNE may play a causal role in driving mesenchymal drift in the RPE, as GSTA4 counteracts these changes (**Figure 3J**). It would be interesting to explore how broadly GSTA4 can reverse mesenchymal drift across tissues and disease contexts.

Unlike the broad effects of partial reprogramming, targeting effectors like GSTA4 offers the potential for a more precise and safer approach to activating rejuvenation. Beyond *in vivo* gene therapy (**Figure 7C**), GSTA4 could be used to enhance iPSC-RPE for cell replacement therapy ^64^. Identifying a single effector also enables pharmacologic targeting: raising intracellular glutathione with its precursors (GlyNAC) reproduced the GSTA4-dependent protection *in vitro*. Although GlyNAC has been linked to extended lifespan and improved mitochondrial function ^58^, our data reveal its specific potential for treating RPE aging and AMD.

While traditional chemical antioxidants (*e.g.* Vitamin C) work by directly neutralizing ROS to delay the accumulation of cellular “trash”, they can disrupt redox balance and interfere with the signaling functions of ROS, and do not repair damage caused by oxidative stress ^6^. In contrast, detoxification through GSTA4 actively clears toxic byproducts (*e.g.* 4-HNE) generated from oxidative damage (**Figure 7B**). Thus, while antioxidants slow the rate of decline, GSTA4-mediated detoxification plays an additional reparative role that not only prevents further cellular injury but also restores redox balance and mitochondrial function that essentially rejuvenates the cell and provides tissue resilience (**Figures 5** and **6**).

It is worth noting that although GSTA4 regulation by OSK is Tet2-independent, our findings do not conflict with prior studies either focusing on stable DNA methylation changes^9,13^, or showing Tet2-dependent effects of OSK in retinal ganglion neurons, such as axon regeneration and vision restoration ^9,14,15,19^ (**Figure 7A**). Indeed, the role of DNA demethylation in retinal ganglion cells has been strongly supported by a more recent study demonstrating that inhibition of DNMT3a, a process that echoes OSK-mediated TET activation, also promotes axon regeneration and restores vision from neuronal injury ^48^. However, oxidative resilience and cell body regeneration represent distinct cellular processes induced by OSK in two different cell types. The distinct mechanisms by which OSK acts in these different contexts underscores the diversity of restorative programs modulated by OSK, and the described Tet2-independent mechanism here may help explain why sometimes extended or cyclic OSK(M) expression yields superior outcomes than seen with transient expression ^14,18,65^.

Using RNA profiling, ATAC-seq and a seq2PRINT approach for precise transcription-factor–binding inference, we demonstrate that OSK dynamically and rapidly regulates GSTA4 (**Figure 4C**-**4F**), consistent with its oxidative-protection pattern (**Figure 2J**). This reversible activation of GSTA4 by ectopic OSK recapitulates its regulation during early embryogenesis (**Figures S4C-S4D**). It suggests an evolutionarily conserved, stem-cell-intrinsic program that guards against oxidative damage at the stem cell stage, yet upon OSK withdrawal, yields to allow ROS-dependent developmental cues. This regulatory pattern may be an example of the antagonistic pleiotropy theory of aging ^66,67^, in which genetic programs that enhance early-life fitness can incur late-life costs. In this case, we speculate the silencing of OSK and GSTA4 around mid-gestation (∼E7.5) is selected for its developmental advantages, but at the same time it may be, at least partly, responsible for the onset of aging damage ^68^.

Beyond supporting the antagonistic-pleiotropy model, our discovery helps unify the epigenetic Information Theory of Aging ^69^ with the Mitochondrial Free-radical Theory of Aging ^70^: age-related epigenetic silencing of GSTA4 appears to be an upstream switch that unleashes ROS accumulation, promoting mitochondrial and cellular dysfunction. The fact that partial reprogramming in aged tissue can re-engage this dormant detoxification pathway to promote oxidative resilience (**Figure 7A**) suggests that there are additional stem cell-intrinsic anti-aging mechanisms ^71^ that could be repurposed for treating age-related diseases.

Looking forward, applying our functional genomic approach of phenotypic validation of partial reprogramming, correlation of transcriptome changes with function, and targeted genetic screens, provides a generalizable blueprint for deconvoluting complex rejuvenation programs. Applying this strategy more broadly could uncover additional effectors that, alone or in combination, recapitulate a fuller spectrum of OSK’s benefits while bypassing the safety concerns. Ultimately, by dissecting and deploying stem-cell-intrinsic anti-aging pathways, we can open up new avenues to harness early-life rejuvenation programs for combating late-life degeneration.

### Limitation of this study

While our work establishes a GSTA4-dependent oxidative resilience pathway downstream of OSK, several limitations merit consideration. First, we confirmed OSK-GSTA4 mediated oxidative protection in RPE and fibroblasts, but the extent to which this axis operates in other tissues, and whether long term GSTA4 overexpression beginning from mid-age can lead to lifespan extension remains to be tested. Second, OSK delivers broad oxidative resistance, whereas GSTA4 was singled out for its potent protection against the mitochondrial toxin NaIO_3_; additional OSK-induced effectors almost certainly contribute to its full protective spectrum. We believe the additional stress resilience factors induced by OSK remain to be discovered, and the comprehensive transcriptomic and ATAC-seq datasets generated in this study will allow further nominations.

## Supporting information

Supplemental Discussion

---

**Figure S1.**
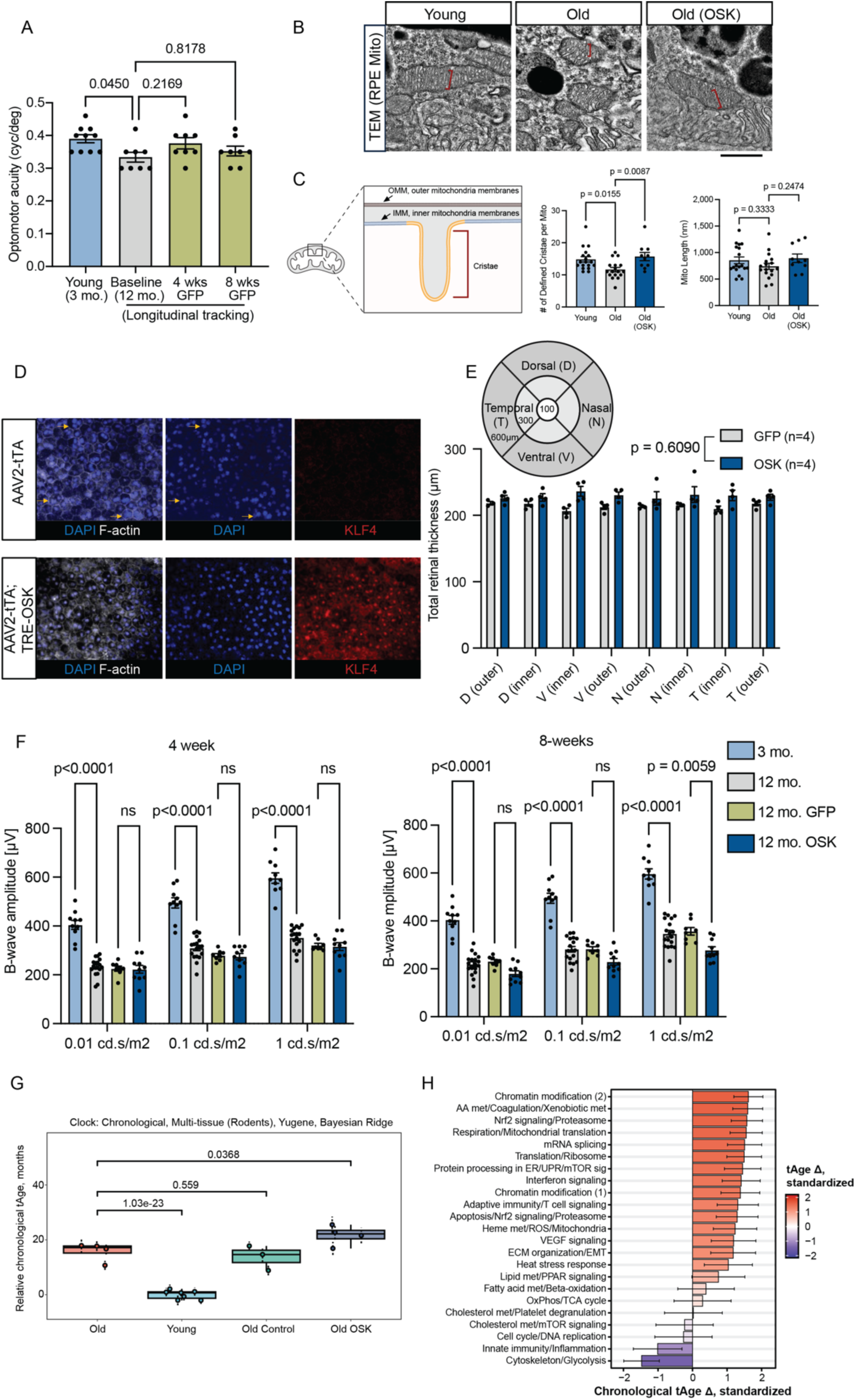
Assessment of the Benefits and Toxicity of AAV2-OSK in Aged RPEs. (A) Quantification of visual acuity by optomotor response (OMR) in 3-month-old mice at baseline, and in 12-month-old mice at baseline, and at 4 and 8 weeks following AAV-GFP injection (4M4F). One-way ANOVA-Bonferroni. (B) Representative high magnification TEM images of RPE mitochondrial cristae in young, old and OSK-treated old eyes. Scale bar, 500nm. (C) Illustration and quantification of defined cristae per mitochondria and mitochondria length in the PRE of young, old and OSK-treated old eyes. (D) Immunofluorescence images of RPE flatmount showing RPE cells in 24-month-old *CFH-H/H* mice maintained on a high-fat, cholesterol-enriched (HFC) diet and treated with AAV-OSK. Multinucleation was defined as ≥3 nuclei per RPE cell. (E) Left: Schematic diagram of mouse retinal subregions. The retina is divided into distinct regions based on radial distance and orientation from the center of the optic nerve head (ONH). Right: Optical coherence tomography (OCT) results showing total retinal thickness and thickness of outer nuclear layer (ONL) in AAV-GFP and AAV-OSK treated eyes (n = 4 eyes per group, equal sex). Two-way ANOVA-Bonferroni. (F) Electroretinogram (ERG) results showing b-wave responses in aged mice at 8 weeks post-OSK injection (Scotopic ERG at strength of 0.01, 0.1 and 1 cd.s/m^2^). Two-way ANOVA-Bonferroni. (G) Transcriptomic age (tAge) of young (4 mo.) and old (17 mo.) mouse RPE cells, untreated and with AAV-OSK treatment, as predicted by rodent multi-tissue transcriptomic clock of chronological age. (H) Standardized chronological tAge differences induced by AAV-OSK treatment in old mouse RPE cells, according to module-specific multi-tissue chronological clocks. Red and blue bars denote pro-aging and anti-aging transcriptomic changes, respectively. Data are mean ± SEM.

**Figure S2.**
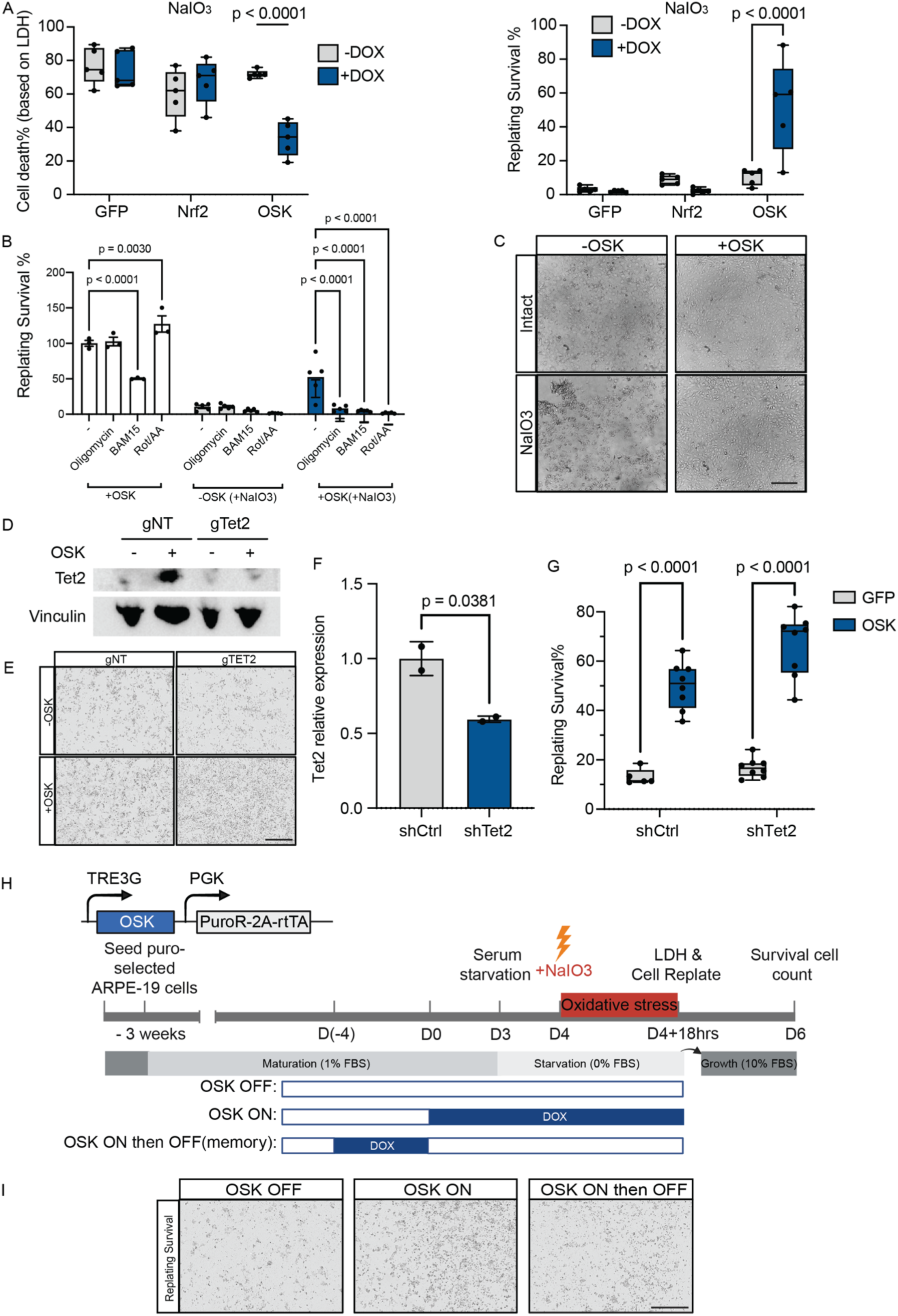
OSK protects RPE from NaIO_3_-induced oxidative stress via a Tet2-independent mechanism. (A) Quantification of NaIO_3_-induced cytotoxicity in matured ARPE-19 cells overexpressing GFP, NRF2, or OSK, assessed by (left) LDH release assay and (right) replating survival assay (n=5 each group). (B) Quantification of ARPE-19 cell survival rates following different mitochondrial inhibitor treatments, with or without OSK induction under normal culture conditions (no NaIO_3_) or oxidative stress conditions (+ NaIO_3_). Oligomycin (blocks ATP production), BAM15 (stops ATP synthesis), and Rot / AA (inhibits Complex I and III) (n ≥ 3). Two-way ANOVA-Bonferroni. (C) Representative brightfield images of mouse fibroblasts derived from OSK-transgenic mice, treated with or without NaIO_3_ in the presence or absence of OSK induction. Scale bar, 200µm. (D) Immunoblot analysis of Tet2 protein levels in ARPE-19 cells that were CRISPR-edited to knockout Tet2 with and without expression of OSK; gTet2, CRISPR-mediated knockout of Tet2; gNT, non-targeting control. Scale bar, 400µm. (E) Representative brightfield images of NaIO_3_ treated ARPE-19 cells that were CRISPR-edited to knockout Tet2 with and without expression of OSK; non-targeting gRNA controls (gNT), Tet2 targeting gRNA (gTet2). (F) qRT-PCR analysis of *Tet2* relative expression in ARPE-19 cells. shTet2, transduction with a short hairpin RNA targeting Tet2; shCtrl, transduction with a non-targeting control short hairpin RNA (n = 2 per group). Two-tailed Student’s t-test. (G) Quantification of cell survival in control (shCtrl) and Tet2-knockdown (shTet2) ARPE-19 cells under NaIO_3_ induced oxidative stress conditions using the cell replating survival assay. The survival of cells with OSK induction (OSK) is compared to control cells expressing GFP (n ≥ 5). Two-way ANOVA-Bonferroni. (H) Schematic of the experimental timeline: matured ARPE-19 cells with doxycycline (DOX)-inducible OSK were exposed to oxidative stress (5 mM NaIO_3_) under different OSK induction patterns and assessed for survival by lactate dehydrogenase (LDH) release and cell replating assays. (I) Representative brightfield images of ARPE-19 cell survival following NaIO_3_ treatment (5mM for 18 hrs). Negative control with OSK off (OFF); positive control with OSK on (ON). RPE cells with OSK on for 4 days followed by withdrawal for 4 days (ON / OFF). Scale bar = 400 µm. All data are presented as mean ± SEM.

**Figure S3.**
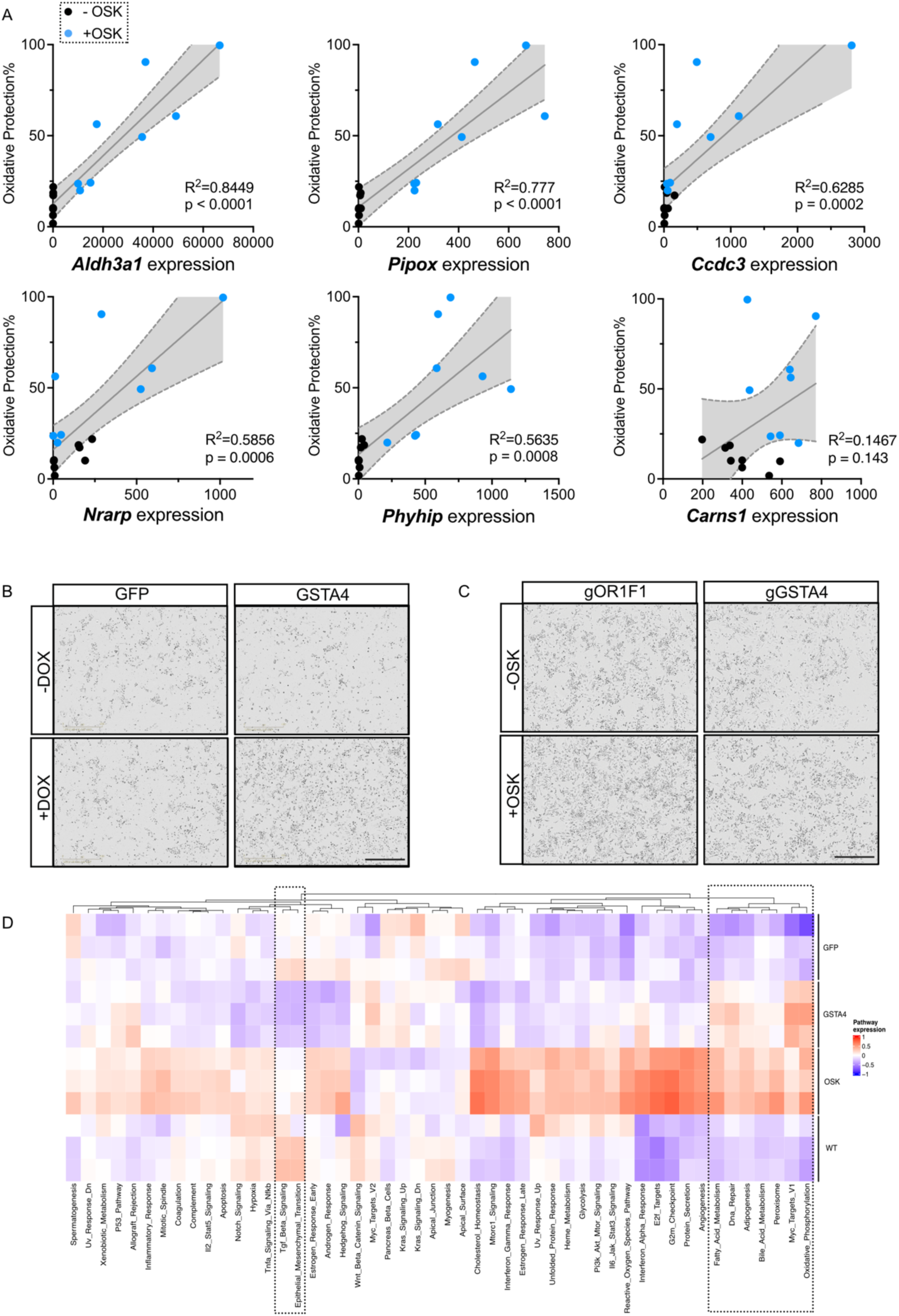
GSTA4 is required for OSK-induced oxidative resilience, and its overexpression partially mimics the transcriptomic effects of OSK. (A) Scatter plots of six other candidate effectors (Left to Right, Top to Bottom: Aldh3a1, Pipox, Ccdc3, Nrarp, Phyhip, Carns1) showing how their mRNA expression correlates with protection (%) against NaIO_3_-induced oxidative stress in mouse fibroblast cells. Black and blue dots represent candidate normalized gene expression without (–OSK) or with (+OSK) induction respectively; solid lines are linear regressions with displayed R² and P values. Shaded regions indicate 95% confidence intervals around the regression line. (B) Representative brightfield images of ARPE-19 stable cell lines with lentiviral integration of inducible GFP or inducible GSTA4, treated with NaIO_3_ in the presence or absence of DOX induction. Scale bar, 400µm. (C) Representative brightfield images of ARPE-19 cells following CRISPR-Cas9 knockout of GSTA4 (gGSTA4) compared to controls (gOR1F1), treated with NaIO_3_ in the presence or absence of OSK induction. Scale bar, 400µm. (D) Gene set enrichment analysis (GSEA) of transcriptomic data from wild type (WT), GFP, OSK, and GSTA4 overexpressing cells (n = 3 per group).

**Figure S4.**
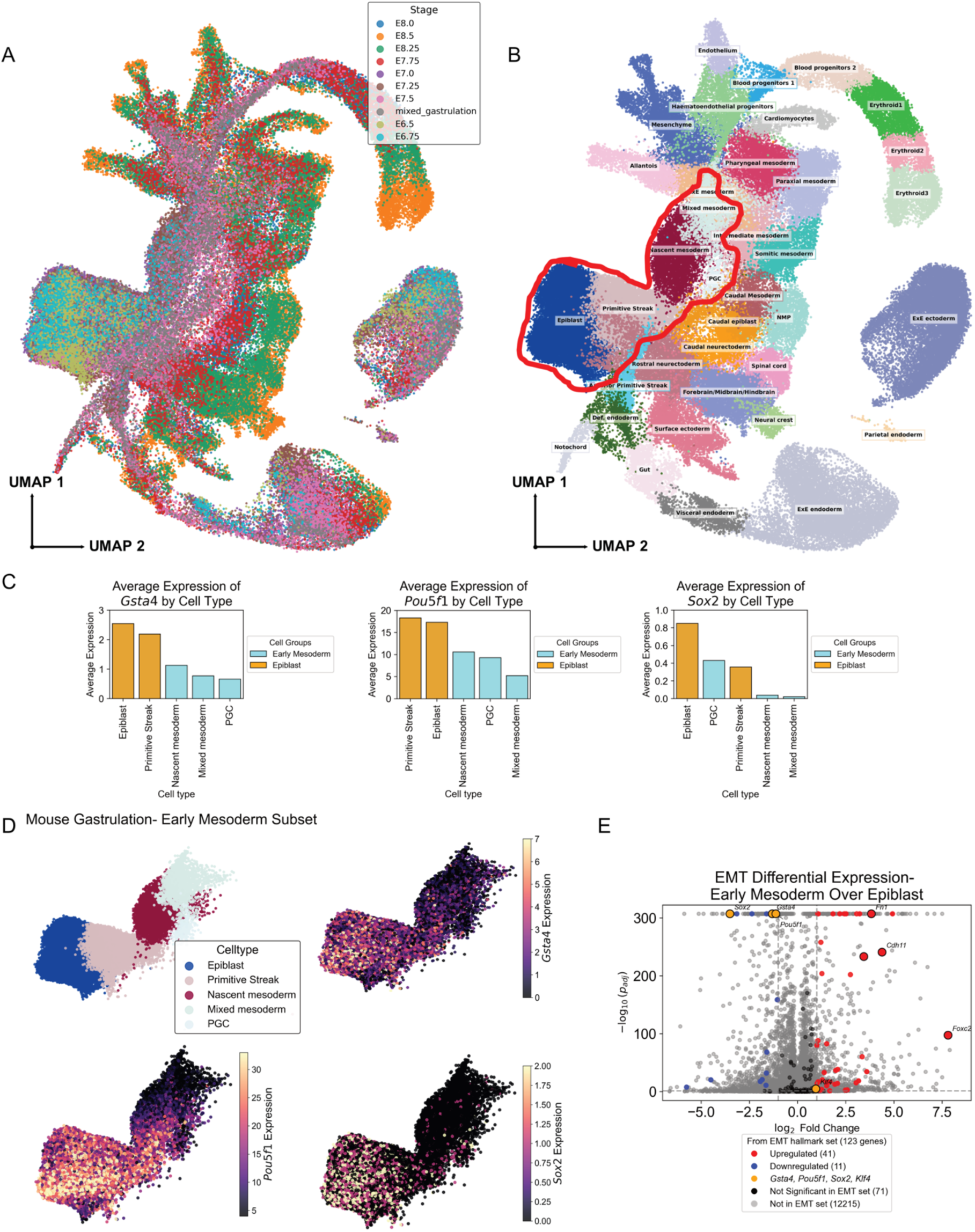
GSTA4, OCT4 and SOX2 are downregulated concurrently with mesoderm specification in early gastrulation. (A) Uniform manifold approximation and projection (UMAP) plot of 116,312 cells from whole mouse embryos spanning the E6.5-E8.5 stages of development. Cells are colored by their developmental time point. (B) Same as (A), but with cells colored by their cell-type annotation. The red highlighted region is the focus of subsequent figure panels. (C) (left to right) Average expression of *Gsta4*, *Pou5f1,* and *Sox2* in each cell type included in the highlighted region from (B). Orange, epiblast-stage cell types; light blue, early mesoderm cell types. (D) Gene expression distribution of Gsta4 across cells of the selected cell subset from the mouse gastrulation dataset, with accompanying expression plots of Sox2 and Pou5f1 shown for reference. Scatter plots of only the cells highlighted in (B), colored by (Upper left) cell-type annotation, (upper right) *Gsta4* expression, (lower left) *Pou5f1* expression, (lower right) *Sox2* expression in the UMAP space. (E) Volcano plot showing the log2-transformed fold change expression (x-axis) and the negative of log10 of adjusted p-value (y-axis) for all genes in the early mesoderm (orange in (C)) over the epiblast (light blue in (C)) cell types. Red, significantly upregulated genes in the EMT hallmark gene set; blue, significantly downregulated genes in the EMT hallmark set; black, EMT genes without significant fold change; orange, *Gsta4, Pou5f1, Sox2, Klf4;* grey, not in the EMT set.

**Figure S5.**
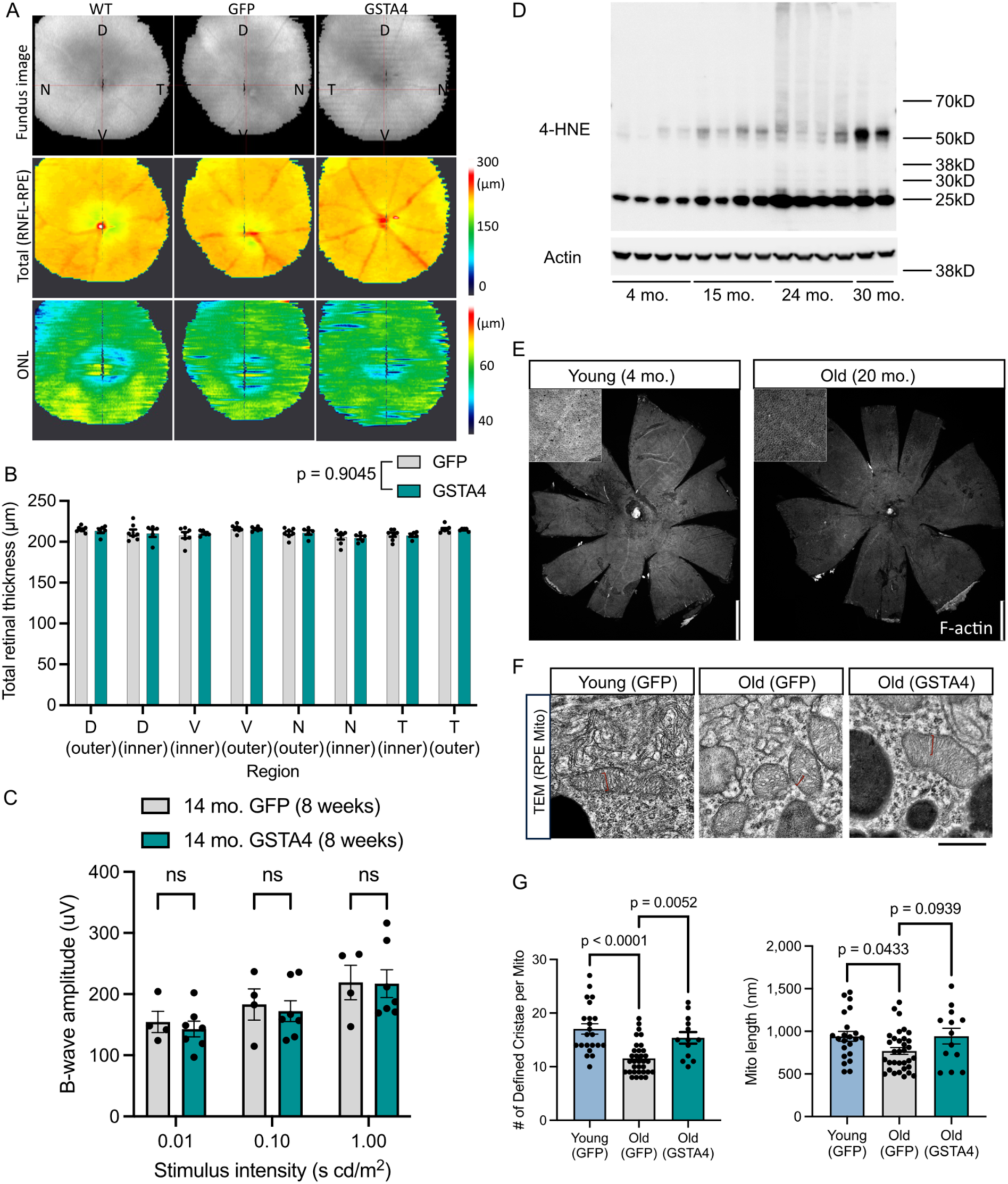
Histological analysis of aged eyes overexpressing GSTA4 demonstrates both safety and efficacy. (A) Comparison of fundus structure, total retinal thickness, and outer nuclear layer (ONL) thickness between untreated (WT) eyes and AAV-treated eyes (GFP group and GSTA4 group) (B) Regional comparison of total retinal thickness measured by OCT between AAV-GFP (n = 7) and AAV-GSTA4 treated eyes (n = 5). Two-way ANOVA-Bonferroni. (C) Electroretinogram (ERG) results showing b-wave responses in aged mice at 8 weeks post-GSTA4 injection (GFP, n = 4; GSTA4, n = 7). (D) Immunoblot analysis of 4-HNE protein conjugates in the mouse retina–RPE tissue lysate across various ages. The major bands are similar in size to those observed in human vitreous fluid^72^. For total protein quantification, the density of each whole lane was measured in Fiji software. (E) Representative immunofluorescence image of RPE flatmounts from young (4-month-old) and old (20-month-old) mice, stained for F-actin (phalloidin). Scale bar, 1 mm. (F) Representative high magnification TEM images of RPE mitochondrial cristae in young, old and GSTA4-treated old eyes. Scale bar, 500nm. (G) Quantification of defined cristae per mitochondria and mitochondria length in the PRE of GFP-treated young, GFP-treated or GSTA4-treated old eyes. All data are presented as mean ± SEM.

**Figure S6.**
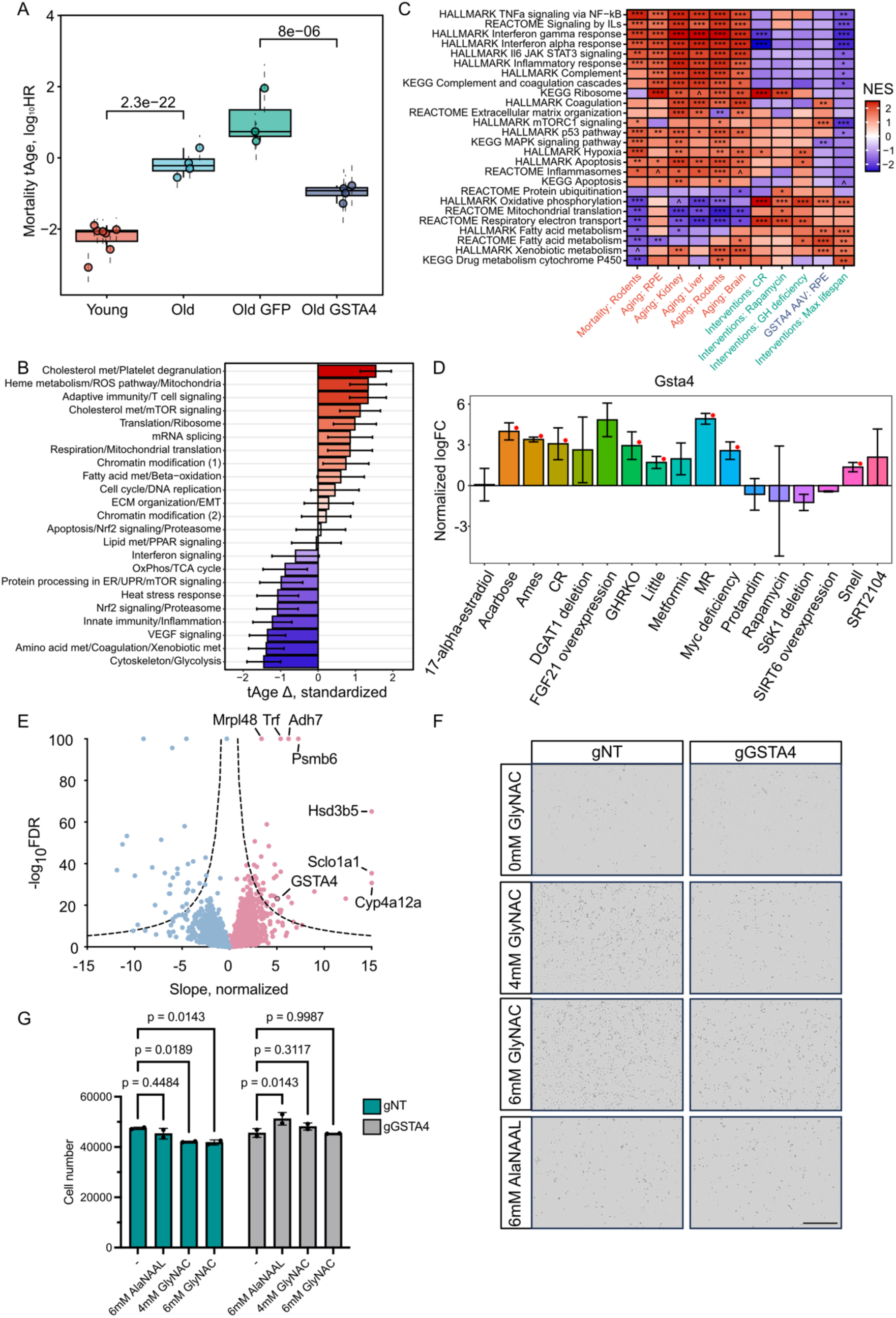
GSTA4 overexpression in RPE restores youthful gene expression, is linked to lifespan-extending treatments, and mediates GlyNAC’s protective effect. (A) Transcriptomic age (tAge) of young (4 mo.) and old (17 mo.) mouse RPE cells, untreated and with AAV treatment, as predicted by multi-tissue transcriptomic clock of expected mortality. (B) Standardized chronological tAge differences induced by AAV-GSTA4 treatment in old mouse RPE cells, according to module-specific multi-tissue chronological clocks. Red and blue bars denote pro-aging and anti-aging transcriptomic changes, respectively. (C) Gene Set Enrichment Analysis (GSEA) results comparing the transcriptomic signatures of GSTA4 AAV with previously identified signatures of mortality, aging, and established longevity interventions. Each cell represents the Normalized Enrichment Score (NES), indicating the degree of enrichment of a given pathway among up- or downregulated genes. HALLMARK, KEGG, and REACTOME ontologies were utilized. (D) Bar chart showing the standardized log fold change of Gsta4 expression in liver across various established lifespan-extending interventions, identified through meta-analysis of transcriptomic data. (E) Volcano blot of genes based on normalized slope of association with maximum lifespan extension in liver induced by longevity interventions. (F) Representative brightfield images of ARPE-19 cells following CRISPR-Cas9 knockout of GSTA4 compared to non-targeting controls (gNT), treated with NaIO_3_ under normal culture conditions and with either 4 mM GlyNAC, 6 mM GlyNAC or 6 mM AlaNAAL. Scale bar, 400µm. (G) Quantification of ARPE-19 cells numbers following CRISPR-Cas9 knockout of GSTA4 (gGSTA4) compared to non-targeting controls (gNT), under no treatment (-) or treatment with either 4 mM GlyNAC (Glycine and N-acetylcysteine), 6 mM GlyNAC or 6 mM AlaNAAL (L-Alanine and N-Acetyl-L-alanine), without oxidative stress conditions (n = 2 per group). Two-way ANOVA-Bonferroni. All data are presented as mean ± SEM. *** p.adj < 0.001; ** p.adj < 0.01; * p.adj < 0.05; ^ p.adj < 0.1.

## EXPERIMENTAL MODEL AND STUDY PARTICIPANT DETAILS

### Data and materials availability

All six RNA-seq datasets and one ATAC-seq dataset generated in this study have been deposited to GEO (GSE304044). Code used for analysis will be available at GitHub. Other data and material generated are available from the corresponding authors upon request.

### Mice

C57BL/6J wild-type mice were housed under 12-h light/dark cycles (6:00/18:00), at an ambient temperature of 70–72 °F (21–22 °C) and 40–50% humidity. All animal procedures were reviewed and approved by the Institutional Animal Care and Use Committees (IACUCs) at Schepens Eye and Ear of Mass Eye and Ear according to appropriate animal welfare regulations. Young and old males and females are obtained from NIA aged rodent colonies.

### Animal Experimental Design

Animal group size was decided using standard deviation estimated from our previous work^9,14^, a two-sided t-test *G*Power* calculation (alpha = 0.05 and 80% power), along with an 20% expected injection failure. Equal numbers of male and female animals were budgeted prior to injection to account for potential sex-dependent effects. Mice with intravitreal bleeding or signs of inflammation (corneal clouding or edema) during or after subretinal injection were excluded based on pre-established criteria. If all injections were successful, additional mice were retained for functional assessments. The examiners of mouse experiments were masked to animal allocation and outcome assessment.

### Subretinal Delivery of AAV2

Subretinal delivery of AAV2 was performed exclusively in the right eye. Following anesthesia, the pupil was dilated using one drop of tropicamide (1%) and phenylephrine (2.5%) ophthalmic solution (PINE Pharmaceuticals, New York, United States). Once adequate dilation was achieved, a 30-gauge needle was used to create a sclerotomy approximately 1 mm posterior to the limbus, between the temporal and superior quadrants. GONAK hypromellose ophthalmic demulcent solution (2.5%) (AKORN, Illinois, United States) was then applied to the corneal surface, and a 5 mm diameter circular contact lens (Warner Instruments, Connecticut, United States) was placed to visualize the fundus under a surgical microscope. A 36-gauge blunt needle (World Precision Instruments, Florida, United States), connected to a 10 μL sub-microliter injection system (World Precision Instruments, Florida, United States), was inserted through the pre-made entry site and guided into the subretinal space, specifically in the nasal-ventral region of the mid-peripheral retina. A total volume of 1 μL of AAV2 (1e12 vg/mL) was delivered into the subretinal space. Successful injection was confirmed by the formation of a visible retinal bleb. Following the injection, residual hypromellose solution was gently removed using an eye spear. To prevent postoperative infection, neomycin and polymyxin B sulfates with bacitracin zinc ophthalmic ointment (Bausch + Lomb, Quebec, Canada) was applied around the injection site.

### AMD CFH Mouse Model

*CFH-H/H* mice were generated and genotyped as described previously^29^. *CFH-H/H* mice were continued on a normal diet (Isopurina 5001; Prolab) and subretinally injected with AAV2-CMV-tTA/TRE-OSK and AAV2-tTA at 120-wk-old. Two weeks post AAV injection, the mice were switched to a HFC diet (TD 88051; Envigo) for 8 wk. All mice were housed conventionally in the same mouse facility under ambient light conditions to control for environmental factors and microbiome fluctuations. Mice were maintained in accordance with the Institutional Animal Care and Use Committee at Duke University.

### Multinucleation Quantification

Mice were perfused with phosphate buffered saline (PBS) before eyes were enucleated. The RPE/choroid/sclera were isolated and fixed overnight in methanol. RPE flatmounts were immunostained with Phalloidin-647 (1:400, Cell Signalling #8940) and nuclei were labeled with DAPI. Confocal images of the flatmounts were collected using Leica TCS SP8 Confocal Microscope. The number of multinucleate RPE cells (≥3 nuclei) per field of view was quantified following the previous study by a masked grader.

### Mice Anesthesia

Mice were anesthetized by intraperitoneal injection of a mixture of ketamine/xylazine (100-200 mg kg−1/20mgkg−1) supplemented by topical application of proparacaine to the ocular surface (0.5%; Bausch & Lomb). For quicker recovery, mice were injected intraperitoneally with yohimbine (2mg/kg) to counteract the anesthesia effects of xylazine, after the procedures.

### Optomotor Response Assessment (OMR)

The visual acuity of mice was evaluated using an automated optomotor reflex-based spatial frequency threshold test with the Optodrum (Striatech). The mice were positioned on a pedestal in the center of an area surrounded by four computer monitors arranged in a quadrangle. These monitors displayed a moving vertical black and white sinusoidal grating pattern. The software captured the mice’s outline, and nose and tail pointers were used to automatically assess their tracking behavior. Tracking behavior was recorded solely when the mice were not moving. The contrast level remained constant at 99.27%, and the rotation speed was set at 12° s−1. The cycle per degree was adjusted using a preprogrammed staircase method. Final thresholds were confirmed by requiring two positive responses, and three negative responses at the next higher spatial frequency. The examiners were masked to group assignments throughout the assessment period.

### Optical Coherence Tomography (OCT)

OCT imaging was performed using a Bioptigen Envisu R-Class OCT system (Leica Microsystems). Mice were anesthetized with a ketamine/xylazine cocktail (100–200/20 mg/kg), and pupils were dilated with 1% tropicamide. Full retinal scans were acquired for all eyes using Bioptigen InVivoVue™ 2.4 software, with the following parameters: 1.4 mm width, 1.4 mm length, 1.79 mm depth, 1000 A-scans per B-scan, 100 B-scans per volume, and 3 frames per B-scan. OCT images were analyzed using Bioptigen InVivoVue Diver 3.4.4 software. Retinas were automatically segmented with the integrated software, and results were reported as whole retinal thickness (ILM to RPE) and outer nuclear layer (ONL) thickness.

### Multifocal Electroretinography (mfERG)

MfERG is a highly localized and sensitive technique for detecting the function of specific areas of the retina, making it particularly valuable when retinal functional improvements are confined to localized regions ^73^. The mfERG methods used in this study have been previously described ^74^. In brief, mfERG tests were conducted using the Celeris pattern stimulator (Diagnosys LLC, Lowell, MA, USA) with Diagnosys mfERG software (v3.8.1) and protocols. Following dark adaptation (≥4 hours), experimental mice underwent pupil dilation (tropicamide 1%) for 15 minutes prior to recording sessions. A 50-degree field of view of the mouse retina was tested using 7 hexagons, each representing a multifocal display area. The pattern ERG stimulator was placed at the center of the cornea, perpendicular to the corneal plane, for each test. Ocular lubrication was maintained using preservative-free carbomer gel 0.18% (Genteal Severe Dry Eye Relief, Novartis Pharma AG, Basel, Switzerland) between corneal surface and the stimulator probe.

### Transmission Electron Microscopy (TEM) Sample Preparation and Analysis

Mice were anesthetized with ketamine/xylazine (100 mg kg−1 and 20 mg kg−1). Live animals were perfused via the aorta with 10 ml of sodium cacodylate buffer (0.1 M, pH 7.4) followed by 10 ml of 1⁄2 Karnovsky’s fixative in 0.1 M sodium cacodylate buffer (Electron Microscopy Sciences). Eyes were enucleated and the anterior segment removed. The eye cups were post-fixed in 2% osmium tetroxide, en bloc stained in 2% aqueous uruanyl acetate, dehydrated and embedded in tEPON-812 epoxy resin. Semithin sections (1 μm) were stained with 1% toluidine blue in 1% sodium tetraborate aqueous solution for light microscopy. Ultrathin sections (80 nm) were cut from each sample block using a Leica EM UC7 ultramicrotome (Leica Microsystems) and stained with 2% aqueous uranyl acetate and Sato’s lead citrate stains using a modified Hiraoka grid staining system. Grids were imaged using an FEI Tecnai G2 Spirit transmission electron microscope (Thermo Fisher at 80 kV interfaced with an AMT XR41 digital CCD camera (Advanced Microscopy Techniques) for digital TIFF file image acquisition. TEM imaging of retina samples was assessed, and digital images were captured at 2k×2k pixel, 16-bit resolution. Mitochondrial quantification methods have been previously described ^75,76^. Briefly, TEM images taken at the same magnification (18,500x) were analyzed using Fiji software. Each mitochondrion and cristae were outlined and measured using the Free Selection tool, while mitochondrial area was quantified using the embedded Region of Interest (ROI) Manager. The number of cristae per mitochondrion was counted. Average measurements in each group were based on data from more than 10 mitochondria across at least 10 cells.

### Corneal Marking for Localization of Subretinal Injection Site in Histological Analysis

Following euthanasia via carbon dioxide inhalation and subsequent cervical dislocation, the nasal-ventral region of the right cornea — approximately 1 mm anterior to the limbus — was cauterized using a high-temperature cautery device with an elongated fine tip (Bovie Medical, Florida, United States). This corneal marking served as a reference for orientation during histological processing. Using curved micro scissors, a triangular notch was created at the marked region, enabling consistent identification of the subretinal injection site. Following fixation, the posterior eye cup was sagittally sectioned through the optic nerve between corneal markings, ensuring both the AAV2-injected and uninjected regions were included within the same histological section.

### Establishment of Stable ARPE-19 Cell Lines with Doxycycline-Inducible Expression

Doxycycline-inducible lentiviral constructs were generated using Gateway LR reactions between pDonor-ORF and pLIX_403 (Addgene #41395). ARPE-19 cells were plated on Day 0 and infected on Day 1 with lentivirus in the presence of 8 µg/mL polybrene. On Day 2, the medium was replaced, and puromycin selection (4 µg/mL) was initiated on Day 4. Selection typically completed by Day 8, as confirmed by complete cell death in uninfected control wells. Stable cells were expanded from Day 9 onward, and plated into wells. Once confluent, cells were cultured in nicotinamide-containing maturation medium for three weeks, with gene expression later induced by 2 µg/mL doxycycline.

### Fibroblast Isolation and Culture

Ear fibroblasts were isolated from Rosa26-M2rtTA/Col1a1-tetOP-OKS mice^9,47^ and cultured at 37 °C in DMEM (Invitrogen) supplemented with non-essential amino acids, 10% tetracycline-free fetal bovine serum (TaKaRa Bio, 631106), and 1% penicillin-streptomycin (ThermoFisher Scientific, 15140122). Fibroblasts were expanded to passage 3 and induced with doxycycline (2 μg/mL) for the indicated durations. All cell lines were confirmed to be mycoplasma-free.

### Seahorse Mito Stress Test

ARPE-19 cells with doxycycline-inducible GFP, OSK, or GSTA4 expression were generated as described above. Cells were plated at 2×10⁴ cells/well in Agilent XFe96 microplates and incubated overnight at 37 °C, 5% CO₂. After three weeks of maturation, doxycycline (2 µg/mL) was added for 48 h, then cultures designated for stress were switched to maturation medium containing 5 mM NaIO_3_ for 24 h to induce mitochondrial oxidative injury. Bioenergetic function was then assessed on the Seahorse XFe96 Analyzer (Agilent Technologies) by measuring real-time OCR and ECAR. Prior to running the assays, the maturation medium was replaced with the assay medium (Seahorse XF Base Medium without Phenol Red, Agilent) supplemented with 2 mM GlutaMAX (ThermoFisher), 1 mM sodium pyruvate (Gibco, Carlsbad, CA, USA), and 25 mM D-glucose (Sigma, St. Louis, MO, USA), adjusted to pH 7.4. Cells were incubated for 1 h in a CO_2_-free 37°C incubator to equilibrate the medium. For the Mito Stress Test, sequential injections of mitochondrial modulators were performed at the following final concentrations: oligomycin (2.5 μM) to inhibit ATP synthase, BAM15 (10 μM) as an uncoupler to assess maximal respiration, and a combination of rotenone and antimycin A (both at 2 μM) to inhibit Complexes I and III respectively. Upon completion of the assays, cells were lysed in ice-cold 1× Cell Lysis Buffer (Cell Signaling Technology, Beverly, MA, USA) supplemented with 1 mM PMSF (Sigma, St. Louis, MO, USA) and stored at -80°C. Protein concentrations were determined using the Pierce BCA Assay Kit (ThermoFisher, Waltham, MA, USA). All metabolic parameters were normalized to the protein content of each well, and data analysis was conducted using the XF Wave software (Agilent Technologies) and exported to GraphPad Prism for statistical analysis.

### CRISPR Knockout

Dual guide RNAs targeting exons of genes of interest were picked from human Brunello library ^77^ (displayed below) and cloned into the lentiCRISPRv2-Opti vector via Golden Gate assembly (NEBridge® Golden Gate Assembly Kit, BsmBI-v2), following the manufacturer’s instructions. Constructed plasmids were transformed into NEB Stable Competent E. coli. Single clones were picked, cultured, and plasmids were isolated via miniprep for sequencing confirmation. For lentivirus production, HEK293T cells were co-transfected with the sequence-verified transfer vector (2 µg) and packaging plasmids pMDL (1.3 µg), pRSV-Rev (0.5 µg) and pMD2.G (VSVG, 0.7 µg) using TransIT transfection reagent. The medium was replaced the next day with ViralBoost Reagent. Viral supernatants were harvested at 48 hours post-transfection, centrifuged at 500g for 5 minutes to remove cell debris and stored at -80°C until use. Target cells were seeded at 2×10^5^ cells per well in 6-well plates 48 hours prior to transduction. For transduction, cells were incubated with 1mL viral supernatant, 8μg/mL polybrene and 1 mL culture medium. The medium was replaced with fresh medium the following day. The transduced cells were passaged into medium containing 4ug/mL puromycin approximately 96 hours post-transduction and selection was maintained for 5 days. Untransduced cells were treated with puromycin as a selection control, and their death confirmed puromycin effectiveness.

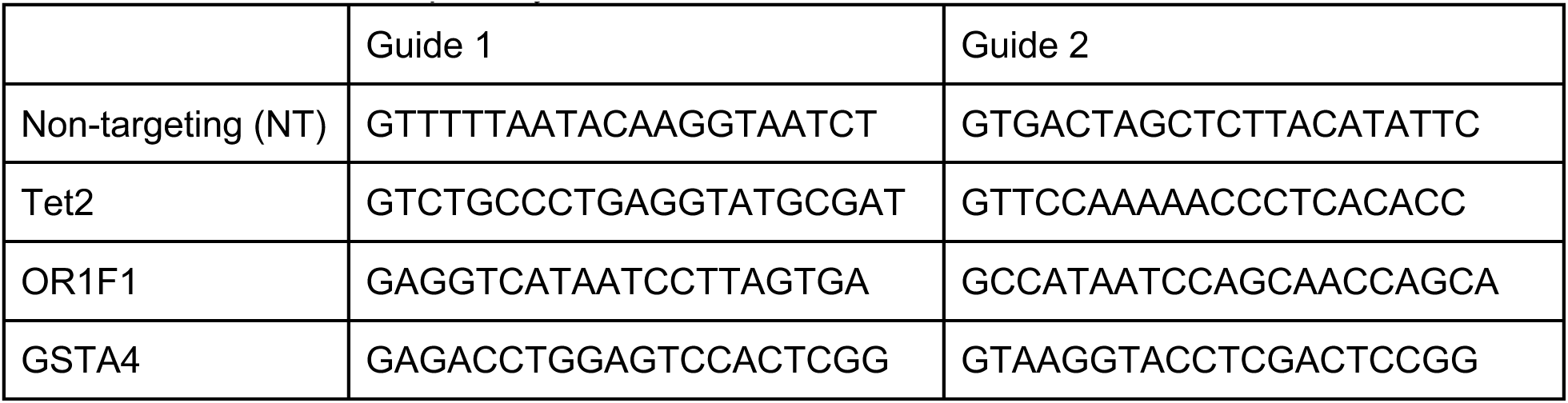

### Replate and Counting

For ORF induction, matured ARPE-19 cells (three weeks in maturation medium) were initially cultured in maturation medium with 2 µg/mL doxycycline for three additional days. Then the cells were switched to starvation medium containing 2 µg/mL doxycycline for an additional day before being treated with different oxidative damage protocols in starvation medium. For each group, one well was left without oxidative damage treatment as a control for counting normalization purposes. Bright field pictures were taken using Incucyte. For LDH assay, 40 µL of cell supernatant was collected and LDH assay reagent was added to the supernatant, followed by incubation at room temperature for 30 minutes under light-protected conditions. The absorbance at 492 nm was measured using a microplate reader after adding LDH assay stop solution. Normal cells treated with lysis solution were used to obtain the maximum LDH release. For replating survival flow count, cells were trypsinized and replated onto a new plate in growth medium and incubated for 1 day. Bright field pictures were taken after washing cells with PBS to remove unattached cells and debris. The harvested cells were then resuspended with 110 µL PBS supplemented with 5% FBS. Samples were analyzed using an Attune Flow Cytometer and a volume of 70 µL per sample was collected.

### RPE Flatmount and Immunofluorescence

Mouse eyes were enucleated and fixed in 4% paraformaldehyde (PFA) at 4 °C overnight. After removing the anterior segment, the neuroretina was gently peeled away from the underlying RPE–choroid cup, which was then radial-cut 7–8 times to generate an RPE flatmount. RPE flatmounts were blocked with BlockAid Blocking solution (Thermo) with 0.1% Triton X-100 for one hour. Then flatmounts were stained overnight with primary antibodies at 4 °C and then secondary antibodies at room temperature for 2 h. Between changes of solutions, all flatmounts were washed 3 times, for 5 min each time. Antibodies used were as follows: goat anti-KLF4 (R&D systems, AF3158), mouse anti-4-HNE (JaICA, MHN-020P), rabbit anti-GSTA4 (Proteintech, 17271-1-AP) and Phalloidin-647 (Thermo, A22287) for F-actin. RPE Flatmounts were mounted with Vectashield Antifade Mounting Medium. KLF4 and GSTA4 antibodies were validated using overexpression cell lines, while the 4-HNE antibody was validated by signal reduction upon GSTA4 overexpression in aged mouse RPEs.

### RPE Flatmount ROS Staining

Retinas were isolated for a whole-mount preparation in cold PBS. Fresh isolated retinas were immersed in a 5-μM concentration of the CellROX Green Reagent (Life Technologies, Rockville, MD, USA) at 4°C for 45 minutes with agitation. After washing with PBS, retinas were fixed with PFA.

### Immunofluorescence Quantification

Immunofluorescence images were acquired using the Leica TCS SP8 confocal microscope (Leica Microsystems). Fluorescence intensity was quantified using ImageJ software. All fluorescence images were processed as 8-bit images in ImageJ to facilitate accurate intensity quantification. The mean gray value of CellROS or 4-HNE staining was measured within predefined regions of interest (ROIs), which were defined using a polygon selection tool in ImageJ. These ROIs corresponded to areas of retinal pigment epithelium (RPE) identified by F-actin staining. OSK- and OSK+ cells were distinguished based on Klf4 staining, while GSTA4- and GSTA4+ cells were identified by GSTA4 staining.

### RPE Pigment Area (%) Quantification

Tissue sections were scanned using a NanoZoomer. After exporting images at 2.5x magnification using NDP.view2, two images at 40x magnification (1432 × 1248 pixels, corresponding to 314.0 × 273.7 μm) were selected from positions approximately 1 mm from the left and right ciliary body, with the left image representing the uninjected site and the right image representing the injected site. Quantification was performed using ImageJ 1.54g / Java 1.8.0_345 (64-bit). Polygon selections were employed to outline the retinal pigment epithelium (RPE) area, while freehand selections were used to delineate the pigment area. The percentage of RPE pigment area (%) was calculated by dividing the pigment area by the total RPE area.

### ATAC-seq Library Preparation and Sequencing

ATAC-seq libraries were prepared using a modified bulk protocol. Briefly, cells were incubated with custom-loaded Tn5 transposase in high-salt transposition buffer for 30 minutes at 37 °C with agitation. Transposed DNA was purified using the Zymo DNA Clean & Concentrator kit. Libraries were pre-amplified for 5 cycles, then subjected to qPCR to determine the additional number of cycles needed, minimizing amplification bias. Final libraries were purified, quality-checked by gel electrophoresis, and quantified. Sequencing was performed on an Illumina platform using paired-end reads, targeting ∼20 million reads per sample.

### ATAC-seq and seq2PRINT Analysis

ATAC-seq data was aligned to the mm10 genome using bowtie2, and duplicate reads were removed using Picard MarkDuplicates. Aligned data was processed using samtools and converted to fragments files with bedtools bamtobed. To visualize the ATAC-seq-based accessibility tracks across conditions, Tn5 insertion was mapped at single base-pair resolutions, normalized by the total sequencing depth of each sample, and then smoothed with a 250 bp running-mean window. High confidence TF binding site annotations were obtained from Unibind (https://unibind.uio.no/)^78^. seq2PRINT ^52^ was used to infer TF binding patterns in individual bulk-ATAC samples. A separate seq2PRINT model was trained for each sample to calculate TF binding scores.

### Mouse RPE Isolation

Eyes were enucleated and placed in cold PBS on ice. The anterior segment was removed, and retinas were either collected separately and snap-frozen or discarded. The posterior eyecup was briefly rinsed in cold PBS, then transferred into 200 µL RNAprotect Cell Reagent (Qiagen) and incubated at room temperature for 10–30 minutes. Tubes were agitated to release RPE cells, and the remaining eyecup was removed. RPE cells were pelleted by centrifugation (2,500 rpm, 5 min, RT), resuspended in 200 µL TRIzol, and flash-frozen on dry ice. RPE purity was confirmed by RNA-seq, showing high expression of Rpe65 and low expression of Cx3cr1 (microglia marker), Cdh5 (endothelial marker), and Rho (photoreceptor marker).

### RNA-seq Library Preparation and Paired-End Sequencing

Total RNA was extracted using TRIzol Reagent (Thermo Fisher) with 1 µL glycogen (10 mg/mL) added to enhance yield. RNA quantity and integrity were assessed using a Qubit 3.0 Fluorometer (Life Technologies) and Agilent TapeStation, respectively. RNA-seq libraries were prepared at Genewiz using the SMART-Seq v4 Ultra Low Input Kit (Clontech) for full-length cDNA synthesis and Nextera XT (Illumina) for library construction. Libraries were indexed via limited-cycle PCR, and quality was confirmed by Qubit and TapeStation. Libraries were multiplexed and clustered on two lanes of an Illumina HiSeq flow cell and sequenced using a 2 × 150 bp paired-end configuration. Image analysis and base calling were performed using HiSeq Control Software, and raw BCL files were converted to FASTQ and demultiplexed using bcl2fastq v2.17, allowing one mismatch for index identification.

### RNA-seq Analysis

Paired-end human and mouse reads were mapped to the canonical chromosomes of the GRCh38/hg38 or GRCm38/mm10 genome genomes, respectively, with STAR (v 2.7.1a)^79^ using a genome index created using Ensembl 96 gene models (with sjdbOverhang=100). Gene counts were obtained for all genes using featureCounts (v1.6.2)^80^ and the same GTF file as for STAR and options “-p -s 1”. Differential expression analysis was performed in R (v 3.6.3) using DESeq2 (v 1.26.0) ^81^ with ‘independentFiltering=FALSE’ and ‘normal’ lfcShrink. Human and mouse expression changes were linked using gene orthologs obtained from Ensembl 110^82^.

### Differential Expression Analysis

Differential expression analysis was performed using DESeq2 with raw RNA counts as input. To calculate the correlation between gene expression and cell survival, raw RNA counts were normalized per sample using the size factor estimated by DESeq2. Log2 fold changes were also estimated using DESeq2.

To calculate the pathway-by-sample gene expression matrix, raw RNA counts were first normalized per sample using the size factor estimated by DESeq2, and then rescaled per gene. The mean scaled expression value of genes in the same pathway was used as the pathway expression score.

### ScRNA-seq Analysis

The mouse gastrulation scRNA-seq dataset was obtained from the Pijuan-Sala, Griffiths & Guibentif *et al*. 2019 study^54^, as indicated following the instructions at https://github.com/MarioniLab/EmbryoTimecourse2018. Cell-type labels provided in the original data object were retained. For **Figure. S4**, the data were subset to cell types “Epiblast”, “Primitive Streak”, “Nascent mesoderm”, “Mixed mesoderm” and “PGC”. The arithmetic mean was used to compute average expression values in each cell type. For **Figure. S4E**, library sizes were normalized to 10 000 transcripts per cell followed by natural-log transformation with a +1 pseudocount. A subset of genes expressed in ≥200 cells was kept. Pairwise differential expression between the epiblast cell types (Epiblast, Primitive Streak) and early mesoderm cell types (Nascent mesoderm, Mixed mesoderm, PGC) was performed with a Wilcoxon rank-sum statistic, with *p-*values adjusted using the Benjamini-Hochberg false discovery rate (FDR-BH) procedure. Gene sets were fetched through **GSEApy v1.1.8**^83^ using the Enrichr interface, focusing on the Hallmark 2020 – **“Epithelial–Mesenchymal Transition”** (EMT) set^84^. Genes with |log₂FC| > 1 and padj < 0.05 were colored as **Up- or Down-regulated** (depending on the sign of the fold change) and visualized using a volcano plot, with a y-axis of −log₁₀ padj plotted against log₂FC and colored as described in **Figure. S4E.**

### Transcriptomic Clock Analysis

Before applying transcriptomic clocks, RPE mouse RNA-seq data underwent filtering and normalization. Genes with fewer than 10 reads in more than 80% of samples were excluded. The filtered data were then processed with Relative Log Expression (RLE) normalization, log-transformation, and YuGene transformation^85^. Missing expression values for clock genes not detected in the dataset were imputed using their corresponding precomputed average values. Normalized gene expression profiles were centered to the median profile of old control samples. Transcriptomic age (tAge) for each sample was estimated using Bayesian Ridge multi-tissue transcriptomic clocks of chronological age and expected mortality^31^. Differences in mean tAge between young and old controls, as well as between old controls and age-matched Gsta4- or OSK-overexpressing samples, were evaluated using a mixed-effects ANOVA model implemented via the rma.uni function from the metafor package in R. Module-specific transcriptomic clocks of chronological age were applied to scaled relative gene expression profiles using the same framework. For each module, tAges were standardized across samples, and then differences in average standardized tAge between old control and treated groups were assessed with ANOVA. Resulting p-values were adjusted for multiple testing using the Benjamini–Hochberg method ^86^.

### Transcriptomic Signature Analysis

To assess association between transcriptomic changes induced by Gsta4 overexpression in RPE cells and signatures of aging, mortality, and lifespan-extending interventions, we performed functional enrichment analysis. The reference signatures included tissue-specific transcriptomic profiles of aging in kidney, liver, and brain, as well as multi-tissue signatures of aging and expected mortality adjusted for chronological age^31,56^. Additionally, we incorporated established hepatic expression signatures of longevity interventions in mice, including those associated with growth hormone deficiency, caloric restriction, and rapamycin, as well as a composite biomarker of maximal lifespan extension derived across dozens of interventions^87^.

For the identification of functional changes induced by aging and Gsta4 overexpression in RPE, we first conducted differential expression analysis using edgeR, comparing (1) young versus old control RPE samples, and (2) old GFP-induced versus old Gsta4-induced RPE samples. Genes were ranked based on a signed log-transformed p-value: -log(pv)×sgn(lfc), where pv and lfc are p-value and logFC of a certain gene, respectively, and sgn is the signum function (equal to 1, -1 and 0 if value is positive, negative or equal to 0, respectively).

Gene set enrichment analysis (GSEA) was then performed on these pre-ranked gene lists using the fgsea package in R (10,000 permutations, multilevel Monte Carlo sampling). Gene sets were derived from the HALLMARK, KEGG, and REACTOME collections in the Molecular Signatures Database (MSigDB). P-values were adjusted for multiple testing using the Benjamini–Hochberg method, and an adjusted p-value < 0.05 was considered statistically significant.

Similarly, GSEA was performed for reference signatures of aging, mortality, and longevity interventions. Enrichment profiles were compared to those observed during RPE aging and in the Gsta4-overexpressing RPE using Spearman correlation of normalized enrichment scores (NES). Association of Gsta4 liver expression change (logFC) across various interventions and its association with intervention effect on mouse lifespan was examined with the mSALT database (https://gladyshevlab.org/mSALT/).

## Acknowledgments

We thank all members of the Weissman lab for helpful discussions. We particularly thank Constance Cepko, Patricia D’Amore, Ya-Chieh Hsu, Kyle Loh, Richard She, Benedikt Brommer, Nasrin Refaian and Xinyao Hu for their technical support and advices, and Michael Shen, Chang Luo, Ming Lei for fellowship support. We are grateful to Caitlin Rausch for assistance with illustrations. We thank Whitehead cores (sequencing, bioinformatics, imaging) for the work. Y.R.L. was supported by postdoctoral fellowships from Glenn/AFAR and Life Sciences Research Foundation, and the NIH Pathway to Independence Award (K99EY037340). S.S. was supported by Bright Focus AMD Fellowship. We thank NIH R01EY031748 (C.B.R), NIH P30EY005722 (to Duke), FFB Free Family AMD Award (to C.B.R.); Research to Prevent Blindness grant (to Duke), P30EY003790 (to Schepens Morphology Core), R01EY03691, P30EYE003790 (to B.R.K. and M.G.K.), R01EY025794, P01AG071463 (to B.R.K). V.N.G was supported by NIA and Hevolution. J.S.W. is an H.H.M.I. investigator. We especially thank Milky Way Research Foundation for the generous support (to J.S.W.).

## Author contributions

Y.R.L., M.M.K., M.G.-K., B.R.K. and J.S.W. conceived the study. Y.R.L led most experiments and data analysis under supervision of B.R.K and J.S.W. Y.R.L., J.C.C., J.A. and Y.H., conducted cell culture, molecular cloning. Y.H., J.C., S.G.J. and G.W.B. assisted with transcriptomic analysis and ATAC-seq. D.Z. and X.Q., assisted with single cell analysis. Y.R.L., H.S., S.S., Z.C., M.M.K., Q.L., P.S., and M.S.-G. conducted mouse experiments, and S.S. developed cornea marking method. A.T., B.Z. and V.N.G. assisted with transcriptomic age analysis. L.C. and C.B.R. conducted *CFH-H/H* mouse experiment. Y.R.L., M.M.K and D.Y.S. performed Seahorse experiments. M.E.-B. assisted with CRISPR knockout experiment. A.M.P., X.T., D.A.S. shared reagents. M.G.-K., M.S.-G., J.D.B. provided advice to experimental design. Y.R.L, B.R.K and J.S.W. co-wrote the manuscript with inputs from co-authors.

