## Supplemental Discussion for "Reprogramming Factors Activate a Non-Canonical Oxidative Resilience Pathway That Can Rejuvenate RPEs and Restore Vision"

Our lab and others have shown that mice display an age-related loss of visual function as measured by a significant reduction in OMR and ERG A, B, and C waves[1–3]. Our current results showed that AAV2-OSK treatment of RPE in aging mice significantly restore their vision from age related decline as assessed by OMR (Figures 1D and S1A) but the OSK treatment did not show a corresponding increase in ERG (Figure S1F). This apparent discrepancy has been observed by other studies indicating discordant OMR and ERG responses occur during different types of retinal degeneration.

In a study of eight transgenic rat lines with different genetic mutations that trigger retinal degeneration, the OKT and ERG responses varied in their ability to detect retinal function changes. The rats that displayed slow retinal degeneration showed a rapid decline in OKT thresholds that did not coincide with ERG measurements[4]. Unexpectedly, rats with rapid retinal degenerations showed slowly declining OKT thresholds that were also inconsistent with ERG measurements[4].

In another study, ERG A-wave and OMR responses were directly compared in a mouse model of retinal oxidative stress induced by increasing concentrations of paraquat delivered by a subretinal injection. At low concentrations of paraquat that induced minimal retinal oxidative damage there was a low correlation between ERG and OMR responses. The correlation between ERG and OMR increased significantly as the concentration of paraquat and the retinal damage increased[5].

From these studies and our own results, we suspect one plausible explanation to our observation is that visual acuity is primarily determined by the functional integrity of high-acuity regions—such as the central retina in humans or the visual streak in rodents—rather than by the average function across the entire retina. Consequently, even localized restoration of RPE health and function can lead to measurable improvements in optomotor response (OMR) scores. In contrast, full-field electroretinography (ERG) reflects the summated activity of the entire retina and is inherently less sensitive to focal changes.

Learning from this, to better assess localized functional gains, particularly those mediated by GSTA4, we acquired a multifocal ERG (mfERG) system, which enables spatially resolved functional measurements and is more appropriate for evaluating region-specific improvements in visual function [6]: mfERG stimulates many retinal areas simultaneously and records localized responses, overcoming full-field ERG's limitation of capturing only a summed signal from the entire retina. Supporting our hypothesis, while there are still no improvements observed in GSTA4-treated eye in the full field ERG (Figure S5C), the mfERG recording indicated enhanced retinal electro-function to light in GSTA4-treated eyes, with higher electrical responses to light stimuli near the AAV injection site (Figures 5C and 5D).

While systematically exploring this with more AAV examples is out of the scope of this study, we encourage colleagues working on AAV-based retinal therapies to assess retinal function using both full-field ERG and multifocal ERG for a more comprehensive evaluation.
